# A single amino acid polymorphism in natural Metchnikowin alleles of *Drosophila* results in systemic immunity and life history tradeoffs

**DOI:** 10.1101/2023.01.16.524277

**Authors:** Jessamyn I. Perlmutter, Joanne R. Chapman, Mason C. Wilkinson, Isaac Nevarez-Saenz, Robert L. Unckless

## Abstract

Antimicrobial peptides (AMPs) are at the interface of interactions between hosts and microbes and are therefore expected to be fast evolving in a coevolutionary arms race with pathogens. In contrast, previous work demonstrated that one AMP, Metchikowin (Mtk), has a single residue that segregates as either proline (P) or arginine (R) in populations of four different *Drosophila* species, some of which diverged more than 10 million years ago. The recurrent finding of this polymorphism regardless of geography or host species, coupled with evidence of balancing selection in *Drosophila* AMPs, suggest there is a distinct functional importance to each allele. The most likely hypotheses involve alleles having specificity to different pathogens or the more potent allele conferring a cost on the host. To assess their functional differences, we created *D. melanogaster* lines with the P allele, R allele, or *Mtk* null mutation using CRISPR/Cas9 genome editing. Here, we report results from experiments assessing the two hypotheses using these lines. In males, testing of systemic immune responses to a repertoire of bacteria and fungi demonstrated that the R allele performs as well or better than the P and null alleles with most infections. With some pathogens, however, females show results in contrast with males where *Mtk* alleles either do not contribute to survival or where the P allele outperforms the R allele. In addition, measurements of life history traits demonstrate that the R allele is more costly in the absence of infection for both sexes. These results provide strong *in vivo* evidence that differential fitness with or without infection and sex-based functional differences in alleles may be adaptive mechanisms of maintaining immune gene polymorphisms in contrast with expectations of rapid evolution. Therefore, a complex interplay of forces including pathogen species and host sex may lead to balancing selection for immune genotypes. Strikingly, this selection may act on even a single amino acid polymorphism in an AMP.

## Introduction

Animals must maintain an intricate balance between their immune reactions to pathogens and host fitness and health^1-5^. This is due to a necessary tradeoff between protecting themselves from numerous encounters with harmful microbes while also avoiding detrimental side effects of their own immune reactions^5-8^. Antimicrobial peptides are at the forefront of animal immune defenses, particularly for invertebrates that rely solely on an innate immune system^9-11^. Invertebrate innate immune defenses against bacteria and fungi are primarily regulated by the immune-deficiency pathway (Imd) (response mainly to gram-negative, some gram-positive bacteria)^12^ and the Toll pathway (response mainly to gram-positive bacteria and fungi)^11,13,14^. When harmful microbes are sensed through various pathogen-associated molecular patterns such as the peptidoglycan in bacterial cell walls, it starts a signaling cascade through the Imd and Toll receptors of host cells^15^. This leads to downstream transcription and translation of molecules including antimicrobial peptides (AMPs)^16,17^. AMPs are generally short, cationic peptides (about 12-50 amino acids) that are often post-translationally processed and secreted from host cells^17,18^. These peptides then directly interact with the pathogens and kill them through mechanisms such as destabilization of membranes or inhibition of essential processes like translation^19,20^. However, AMPs and other immune genes are regulated temporally and spatially to avoid harm to the host, as expression of immune molecules can be detrimental due to excessive inflammation, off-target effects, the energetic cost of AMP expression, or other factors^5,21-26^.

Although population-level immune gene variation can be critical for keeping up with an ever-adapting suite of pathogens^27,28^, very little is known about the functional differences between AMP alleles in terms of how they interact differentially with both the pathogen and the host^29^. Previous work with *Drosophila* has indicated that multiple AMPs are commonly present in multiple allelic forms across different species^29,30^. Indeed, functional work on the Imd-regulated Diptericin A peptide indicates that one allele exhibits significantly stronger antimicrobial activity against a *Providencia rettgeri* bacterial pathogen challenge^29^. This would indicate that some alleles may be associated with higher host fitness in the presence of specific pathogen infections. Further, AMPs in *Drosophila melanogaster* and *Drosophila mauritiana* show evidence of balancing selection based on calculations of Tajima’s D, π, and Watterson’s θ^31^. These are three population genetic measures that collectively indicate higher observed levels of nucleotide diversity than expected in several populations. Observed AMP nucleotide diversity is also higher than observed across the genome broadly or across all immune genes^31^. These results are consistent with maintenance of AMP variation through balancing selection. Maintenance of alleles is in contrast with expectations of rapid evolution of AMPs due to direct interactions with pathogens and an anticipated evolutionary arms race between AMPs and microbes^32-36^. This introduces the question of how and why AMP alleles are balanced in *Drosophila* populations.

One intriguing AMP is Metchnikowin (Mtk), a 26-residue AMP whose expression is controlled by both the Toll and Imd pathways^37,38^. It exhibits activity against a variety of fungi, bacteria, and potentially eukaryotic parasites^14,38-40^. Mtk is expressed in various tissues in response to infection including the fat body (comparable to human liver), surface epithelia, and the gut^41,42^. It is also a proline-rich AMP, others of which are known to inhibit pathogen ribosomes^43^. Functional research in a heterologous yeast system indicates it may interact with pathogen succinate-coenzyme Q reductase or possibly critical fungal cell wall components^44,45^, which may be pathogen targets. Further work suggests that Mtk may play a role in the fly nervous system^46^. Flies lacking the gene exhibit improved outcomes after traumatic brain injury, indicating that Mtk may have additional functions in the host beyond pathogen defense^46^. Further, Mtk may play important roles in sleep and memory functions^47^, as well as post-mating responses in females^48^. Importantly, previous work identified two alleles of *Mtk* that are likely balanced in various populations of *D. melanogaster, D. simulans, D. mauritiana*, and *D. yakuba*^30^. In these populations, the mature peptide has either a proline (MtkP) or an arginine (MtkR) in the third to last amino acid position. This is despite the fact that species divergence time ranges from an estimated 3-12 million years ago^49,50^. The probability of these same two alleles being found in all four species by random chance is therefore exceedingly low. This indicates a high likelihood that these alleles each play functionally important roles in these populations. Two main hypotheses could explain this repeated functional balance between the two alleles^31^. The *autoimmune hypothesis* poses that one of the alleles has more potent antimicrobial activity, but carries a negative fitness effect in the host in the absence of infection. The expectation with this hypothesis is that flies with this allele would have higher survival with infection, but reduced relative fitness in the absence of infection. The *specificity hypothesis* poses that each allele is more potent against a different suite of pathogens. The expectation with this hypothesis is that flies with each allele would have higher survival depending on the specific pathogen, but neither would necessarily have a higher fitness cost without infection.

Here, we use CRISPR/Cas9-edited *Drosophila melanogaster* strains with either *MtkP, MtkR*, or *MtkNull* alleles in various life history and infection assays to assess these hypotheses and expectations. We find that in males, the evidence broadly supports the autoimmune hypothesis, where flies with the *MtkR* allele have lower fitness without infection, but generally survive just as well or better than the other alleles with infection. However, female infections show some contrasting results, where either allele can have stronger antimicrobial activity depending on the pathogen. When considering sex by genotype interactions, the evidence therefore also supports the specificity hypothesis, suggesting that both models may contribute to maintenance of the alleles in fly populations. This is one of the first cases where functional work consistently and clearly demonstrates that even a single amino acid difference between AMPs may contribute significantly to host immune function and help maintain population-level diversity in AMPs.

## Results

### Fly lines with CRISPR/Cas9-edited Metchnikowin alleles were generated

To assess our two hypotheses, we generated CRISPR/Cas9-edited *D. melanogaster* that are genetically identical except for their *MtkP, MtkR*, or *MtkNull* alleles. We created multiple strains of each, with three independent isolines derived from unique CRISPR-edited individuals for each allele type (Figure S1). The three *MtkP24* (henceforth, *MtkP*) alleles were derived from siblings of the founders of edited lines, ensuring that they were exposed to the same injection history as the edited lines. This allowed experimentation on three independent *MtkP* strains, three independent *MtkP24R* (henceforth, *MtkR*) strains, and three independent *MtkNull* strains. Notably, the *MtkNull* strains each have unique deletions. One has a single base pair (bp) deletion, another a 6 bp deletion, and a third has a 17 bp deletion. Two of these (1bp and 17bp) result in frameshifts, and the other removes two amino acids. Experiments and analyses were performed using three independent strains of each allele, and statistics were employed in all analyses to assess any potential differences among the three strains of an allele type (Table S1). It was broadly the case where phenotypically and statistically, the isolines grouped together by allele type in the following experiments. Between this approach and sequencing confirming no off-target effects of CRISPR/Cas9 (Figure S1, Table S3), results are robustly supported and reduce the likelihood of any results being impacted by other genetic elements.

### *MtkR* allele flies exhibit lower egg hatch rates and die earlier as adults

We performed a life history assay with the *Mtk* strains to assess any fitness differences as anticipated by the autoimmune hypothesis. This assay involved tracking the offspring of dozens of individual male and female pairs from eggs to the end of adulthood (Figure 1a). This allowed quantification of offspring survival between stages for many families. While offspring numbers, survival rates between developmental stages, and the proportion of females were similar or exhibited small differences from allele to allele (Figure S2, Table S1), there were a few notable life history differences (controlling for eggs laid and vial differences). The first was the egg-to-larval hatch rate, where approximately 5% fewer *MtkR* eggs hatched compared to *MtkP* or *MtkNull* (Figure 1b, logistic regression & Tukey contrasts *MtkR* vs *MtkP* or *MtkNull*, ***p<1×10^−5^). The second was adult longevity, where *MtkR* flies died slightly but significantly earlier than their counterparts (Figure 1c, ANOVA, ***p_Genotype_<2×10^−16^). The difference in longevity was larger in females. In addition, male death rates with *MtkR* vary from experiment to experiment, but earlier female death remains consistent (Figure S3, ANOVA, Block A ***p_GenotypexSex_=4.25×10^−7^, Block B ***p_GenotypexSex_=1.28×10^−6^). Third, *MtkNull* males exhibited improved longevity over both alleles. Overall, these results suggest that *MtkR* carries a slight fitness cost in the absence of infection, as expected under the autoimmune hypothesis.

**Figure 1.**
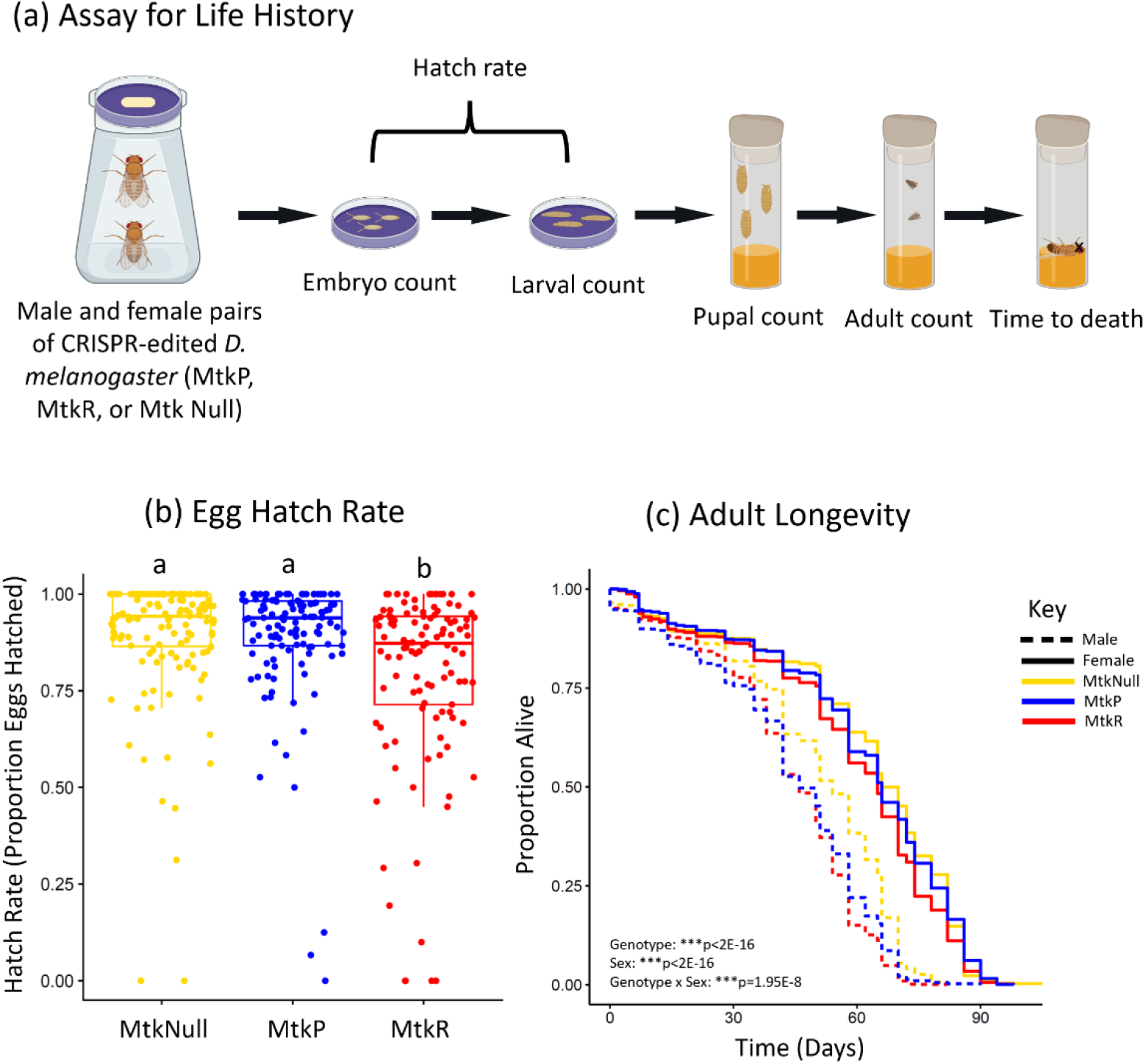
*MtkR* allele flies have lower egg hatch rates and adult longevity times. (a) Overview of the experimental design, where individual pairs of males and females of a given genotype were crossed. They were allowed to lay eggs for 24 hours and these embryos were counted. They were then monitored for hatching rates and the larvae were moved into fly food vials. The pupae were counted each day, along with adult male and female emergence, and these adults were tracked over time until death. (b) The egg-to-larvae hatch rate was counted for each family. Each dot represents the offspring of a single male and female of the indicated genotype. Each dot represents the proportion of all eggs that hatched from one dish (mean eggs per dish was 41). The boxes indicate the interquartile range. Outer edges of the box indicate 25^th^ (lower) and 75^th^ (upper) percentiles and the middle line indicates 50^th^ percentile (median). Whiskers represent maximum and minimum ranges of data within 1.5 times the interquartile range of the box. Letters indicate statistical significance groups, based on a logistic regression and Tukey post hoc test. (c) Lines represent adult longevity beginning at emergence from pupae. Each genotype contains an average of 3879 total flies (males plus females). Statistics based on an ANOVA with Tukey post-hoc test (Table S1). The entire experiment was performed twice, and graphs represent a combination of data from both experiments.

### Male *MtkR* allele flies survive systemic fungal infection as well or better than *MtkP* allele or *MtkNull* flies

We performed a series of systemic infections in flies by piercing a pathogen-dipped needle into the thorax and measuring survival each day over time to determine whether the different *Mtk* alleles provided different levels of protection against pathogens. We first performed fungal infections in males of the different alleles and measured survival over 21 days (Figure 2, Figure S4). We also included 2 additional fly lines that are deficient for either the *spz* or *myd88* genes to compare loss of *Mtk* to loss of the entire Toll pathway and ensure that infections were resulting in expected levels of death for Toll-deficient lines. However, note that these lines are on different genetic backgrounds and so are not as comparable but give a larger scale view. Results revealed that *Mtk* alleles play important roles in various infections in males. First, *Mtk* is important for survival after infection with a variety of filamentous fungi and yeast including *Fusarium oxysporum, Aspergillus fumigatus, Aspergillus flavus, Candida glabrata*, and *Galactomyces pseudocandidus*. This result is based on improved survival of flies with either one or both *Mtk* alleles over *MtkNull* flies. Second, in every case in Figure 2, the *MtkR* allele performed as well or better than the *MtkP* allele. Importantly, survival of *MtkR* alleles with *F. oxysporum, B. bassiana*, and *G. pseudocandidus* was significantly higher than those with the *MtkP* allele (Figures 2a,b,f, logistic regression & Tukey correction *MtkP* vs *MtkR, F. oxysporum* ***p<0.001, *B. bassiana* ***p<0.001, *G. pseudocandidus* **p=4.56×10^−3^), suggesting that there is a pattern in at least some cases where *MtkR* is the more potent allele in *vivo*. Notably in the case of *B. bassiana* infection (Figure 2b), the *MtkP* allele was associated with an even lower survival than the *MtkNull*, suggesting an unexpected negative interaction between this allele and this pathogen (logistic regression & Tukey correction *MtkP* vs *MtkNull*, ***p<0.001). These results, in addition to previous results showing a fitness cost with no infection (Figure 1), continue to support the expectations of the autoimmune hypothesis. Finally, for several fungal infections (including *Candida albicans*), we did not find significant differences in allelic survival (Figure S5a).

**Figure 2.**
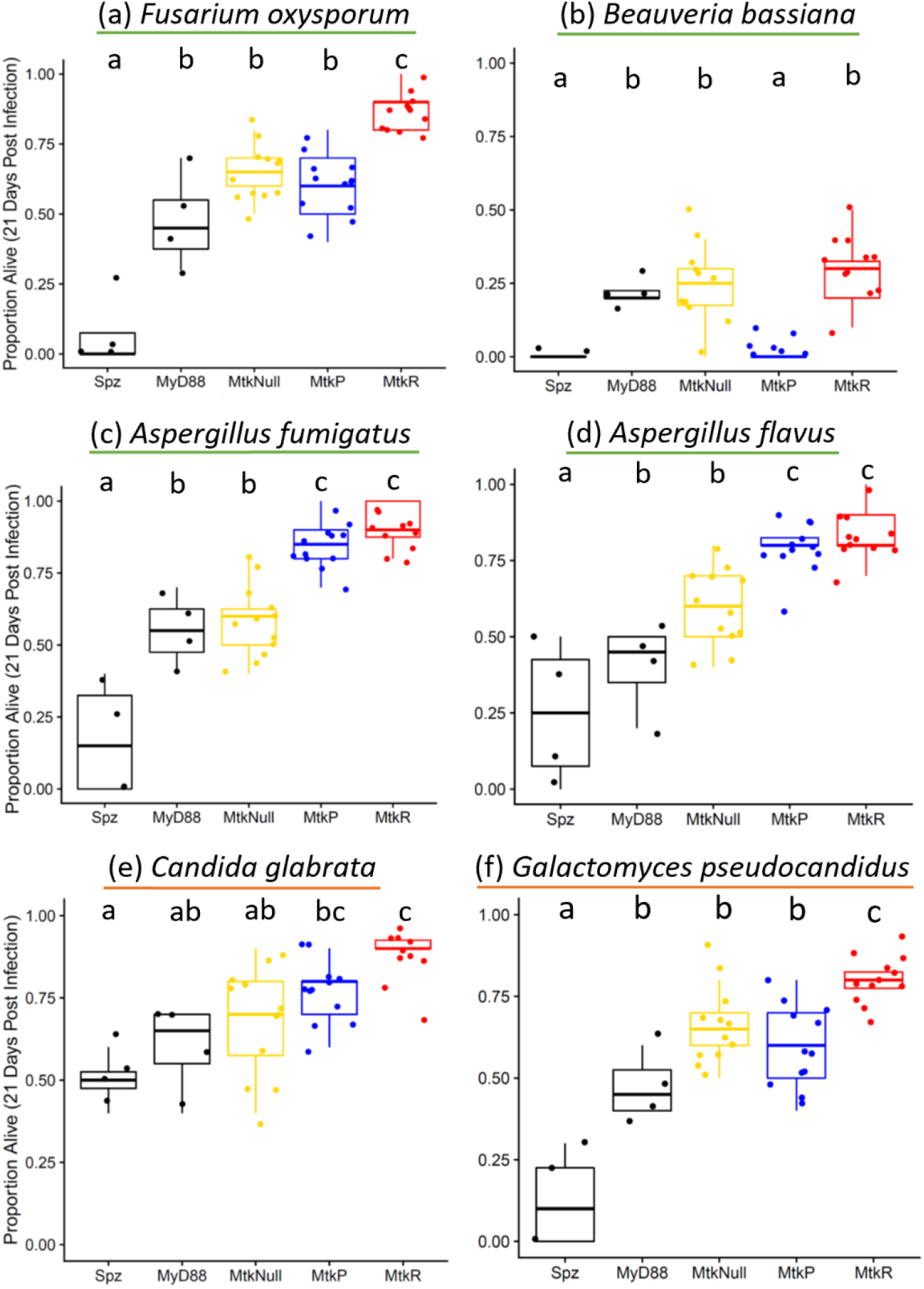
*MtkR* male flies perform as well or better than other genotypes against a variety of filamentous fungal and yeast infections. Infections were performed with the indicated microbes, using either spores (green underline) or yeast cultures (orange underline). Each dot represents the survival 21 days after infection for a vial starting with 10 males. Each set of data represents two independent experiments combined. Corresponding survival curves and controls for this experiment are shown in Figure S4. The boxes indicate the interquartile range. Outer edges of the box indicate 25^th^ (lower) and 75^th^ (upper) percentiles and the middle line indicates 50^th^ percentile (median). Whiskers represent maximum and minimum ranges of data within 1.5 times the interquartile range of the box. Letters indicate statistical significance groups, based on a logistic regression and Tukey post hoc test (Table S1).

### *Mtk* allelic variation is associated with differential survival after infection with some bacterial pathogens

Metchnikowin is canonically expressed in response to both fungal and some bacterial pathogens, and was initially characterized with potent *in vitro* activity against both *Micrococcus luteus* and *Neurospora crassa*^38^. To determine whether *Mtk* genotype is associated with differential ability to survive bacterial infection, we performed additional systemic infections using a variety of Gram-positive bacteria (*Bacillus thuringiensis, Enterococcus faecalis, Lysinibacillus fusiformis, Staphylococcus succinus, Staphylococcus sciuri, Lactococcus brevis*, and *Lactococcus plantarum*), since the Toll pathway primarily controlling *Mtk* largely responds to Gram-positive bacteria and fungi. We also tested two Gram-negative bacteria (*Providencia rettgeri* and *Serratia marcescens*) (Figure 3, Figure S5b-f, Figure S6). We find that while many infections result in no differences in survival across alleles (Figure 3c, Figure 3d, Figure S6), both *B. thuringiensis* and *P. rettgeri* infections do (Figure 3a, Figure 3b). The *MtkR* flies survive at higher rates compared to *MtkP* flies (logistic regression & Tukey correction *MtkR* vs *MtkP, B. thuringiensis* *p=4.45×10^−2^, *P. rettgeri* *p=3.12×10^−2^). Notably and similar to the results with *B. bassiana* infection, both of these bacterial infections resulted in reduced survival of the *MtkP* allele compared to *MtkNull*. This result again suggests there is a negative interaction between the *MtkP* allele specifically with certain pathogen infections.

**Figure 3.**
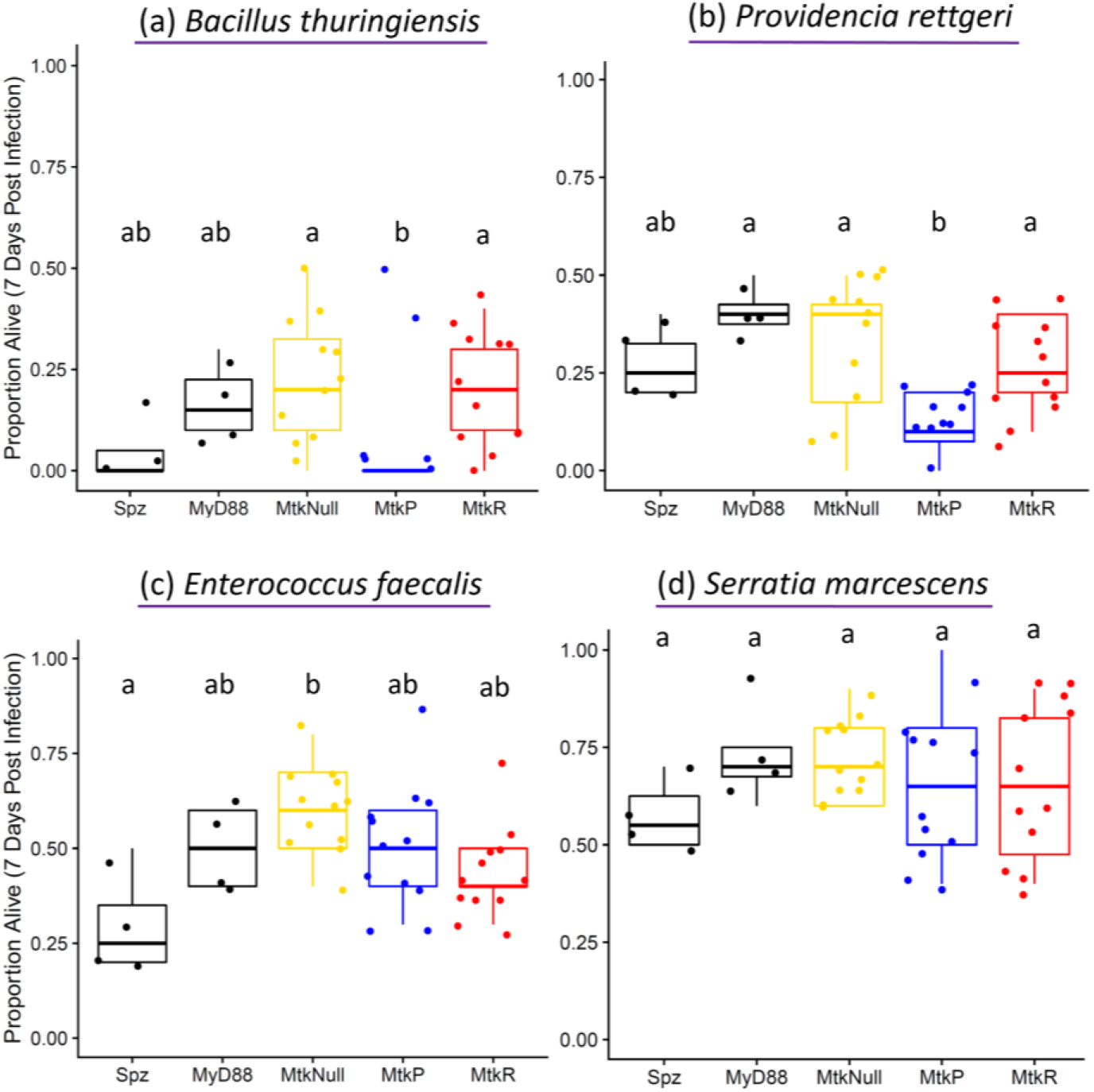
*Mtk* alleles are associated with differential survival after infection with some bacterial infections. Infections were performed with the indicated microbes, using bacteria (purple underline). (a) and (c) are Gram-positive, and (b) and (d) are Gram-negative). Each dot represents the survival 7 days after infection for a vial starting with 10 males. Each set of data represents two independent experiments combined. Corresponding survival curves and controls for this experiment are shown in Figure S6. The boxes indicate the interquartile range. Outer edges of the box indicate 25^th^ (lower) and 75^th^ (upper) percentiles and the middle line indicates 50^th^ percentile (median). Whiskers represent maximum and minimum ranges of data within 1.5 times the interquartile range of the box. Letters indicate statistical significance groups, based on a logistic regression and Tukey post hoc test (Table S1).

### There is an interaction between sex and *Mtk* allele for survival after infection

Since there are commonly differences between male and female innate immune functions^51-55^, we also performed systemic infections with females and a subset of representative bacterial and fungal pathogens (Figure 4, S7). One bacterial species with male allelic differences (*P. rettgeri*) and one without (*E. faecalis*), one sporulating fungus with allelic differences (*B. bassiana*), and one with similar allelic performance (*A. fumigatus*), and one yeast (*C. glabrata*) were chosen. Collectively, these represent the full range of phenotypes in male infections to compare with females. Results demonstrated several key similarities and differences compared to male infections. First, all lines had similar survival rates in *E. faecalis* infection but *MtkR* improves survival with *C. glabrata* infection, similar to males (logistic regression & Tukey correction *MtkR* vs *MtkNull*, *p=2.0×10^−2^). However, in contrast with males, *Mtk* alleles are not associated with survival after infection with *P. rettgeri* or *A. fumigatus* in females. Notably, female infection with *B. bassiana* also differed from males where the *MtkP* allele females survived at higher rates than *MtkR* flies while the opposite is true for males (logistic regression & Tukey correction *MtkP* vs *MtkR*, **p=2.33×10^−3^). To do a more direct side-by-side comparison of male and female differences, we performed additional infections with both sexes using *A. fumigatus* and *B. bassiana* as representative pathogens of the two differing phenotypes (important in males but not females for *A. fumigatus*, opposite survival of alleles for *B. bassiana*) (Figure 5, S8). This confirmed the differences, where *Mtk* alleles are important in male *A. fumigatus* infection but not female (ANOVA *MtkP* & *MtkR* only, ***p_Sex_=8.31×10^−15^), and where the *MtkP* allele is beneficial for females and the *MtkR* allele is beneficial for males in *B. bassiana* infection (ANOVA *MtkP* & *MtkR* only, ***p_GenotypexSex_=5.2×10^−8^).

**Figure 4.**
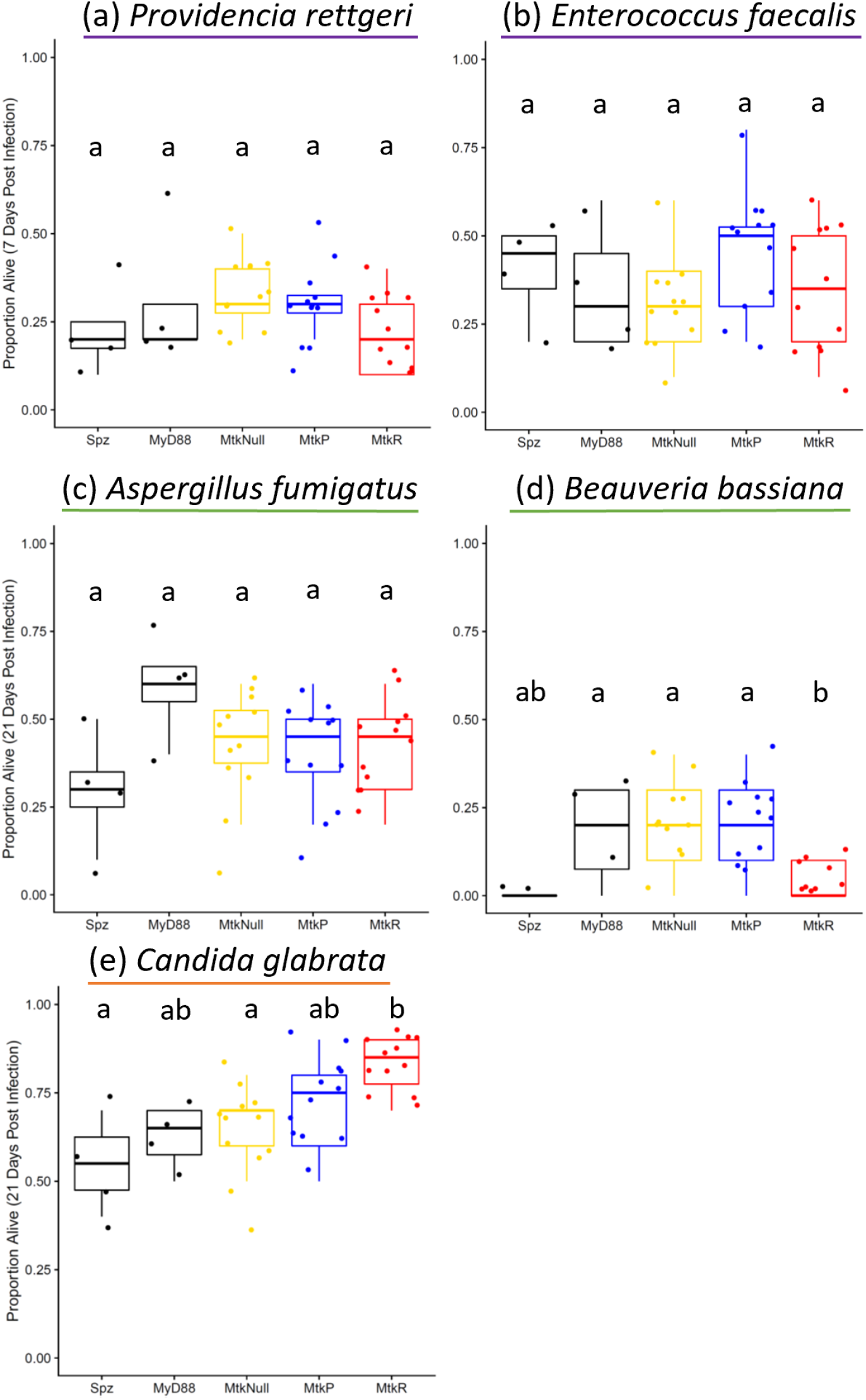
Survival against some infections depends on host sex along with *Mtk* allele and pathogen species. Infections were performed in females with the indicated microbes, using bacteria (purple underline), spores (green underline), or yeast (orange underline). Each dot represents the survival 21 days after infection for a vial starting with 10 females. Each set of data represents two independent experiments combined. Corresponding survival curves and controls for this experiment are shown in Figure S7. The boxes indicate the interquartile range. Outer edges of the box indicate 25^th^ (lower) and 75^th^ (upper) percentiles and the middle line indicates 50^th^ percentile (median). Whiskers represent maximum and minimum ranges of data within 1.5 times the interquartile range of the box. Letters indicate statistical significance groups, based on a logistic regression and Tukey post hoc test (Table S1).

**Figure 5.**
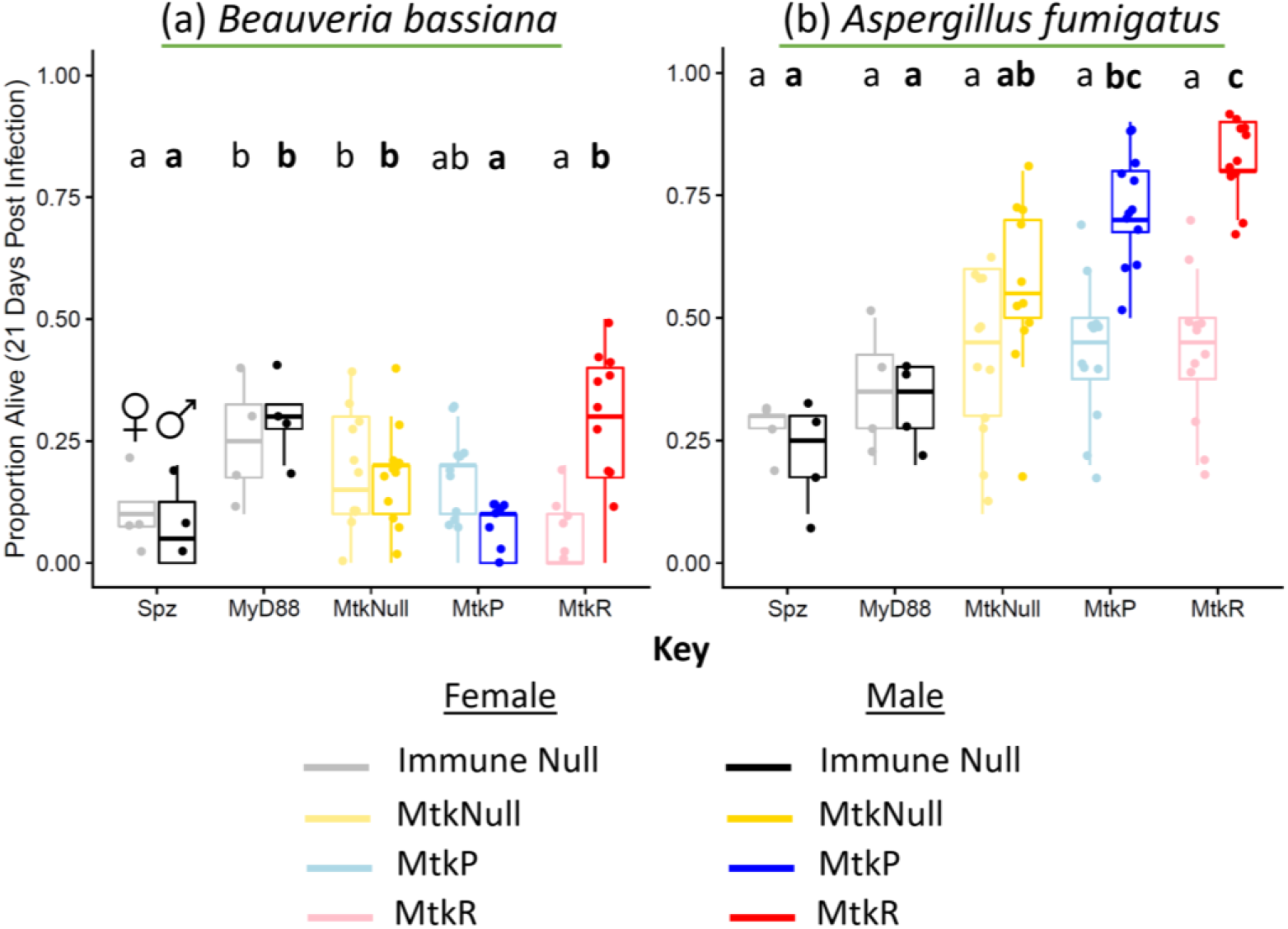
Side-by-side male and female fungal spore infections demonstrate key differences in phenotypes. Infections were performed in females and males with the indicated microbes, using two representative sporulating fungal species (green underline). Each dot represents the survival 21 days after infection for a vial starting with 10 flies. Each set of data represents two independent experiments combined. Females are shown in the lighter colors on the left of each genotype pair, and males are shown in the darker colors on the right of each genotype pair. Corresponding survival curves and controls for this experiment are shown in Figure S8. The boxes indicate the interquartile range. Outer edges of the box indicate 25^th^ (lower) and 75^th^ (upper) percentiles and the middle line indicates 50^th^ percentile (median). Whiskers represent maximum and minimum ranges of data within 1.5 times the interquartile range of the box. Letters indicate statistical significance groups, based on logistic regression and Tukey post-hoc test (Table S1). Non-bolded letters indicate the females, and bolded letters indicate the males.

### Flies carrying the *MtkR* allele are more resistant to infection than *MtkP*

To assess whether allelic differences in survival are based on pathogen tolerance or resistance^56^, we measured microbial load in males infected with a subset of bacteria and yeast that exhibited no allelic effects (*E. faecalis*), higher *MtkR* survival (*C. glabrata*), or *MtkP* survival below that of *MtkNull* (*P. rettgeri*) (Figure 6). While there were no pathogen load differences as expected in *E. faecalis* infection, *MtkR* had a significantly lower pathogen load for *C. glabrata* (logistic regression & Tukey correction, *MtkNull* & *MtkR* *p=3.2×10^−2^, *MtkP* & *MtkR* **p=2.34×10^−3^) and a lower microbial load for *MtkR* vs *MtkNull* in *P. rettgeri* (logistic regression & Tukey correction, **p=4.3×10^−3^). Also, despite *MtkP* exhibiting lower survival in males with *P. rettgeri*, the load was not significantly different from *MtkNull* flies. These results suggest that *MtkR* survival may be improved by greater pathogen resistance, but higher pathogen loads do not underlie lower survival with *MtkP* vs *MtkNull* alleles.

**Figure 6.**
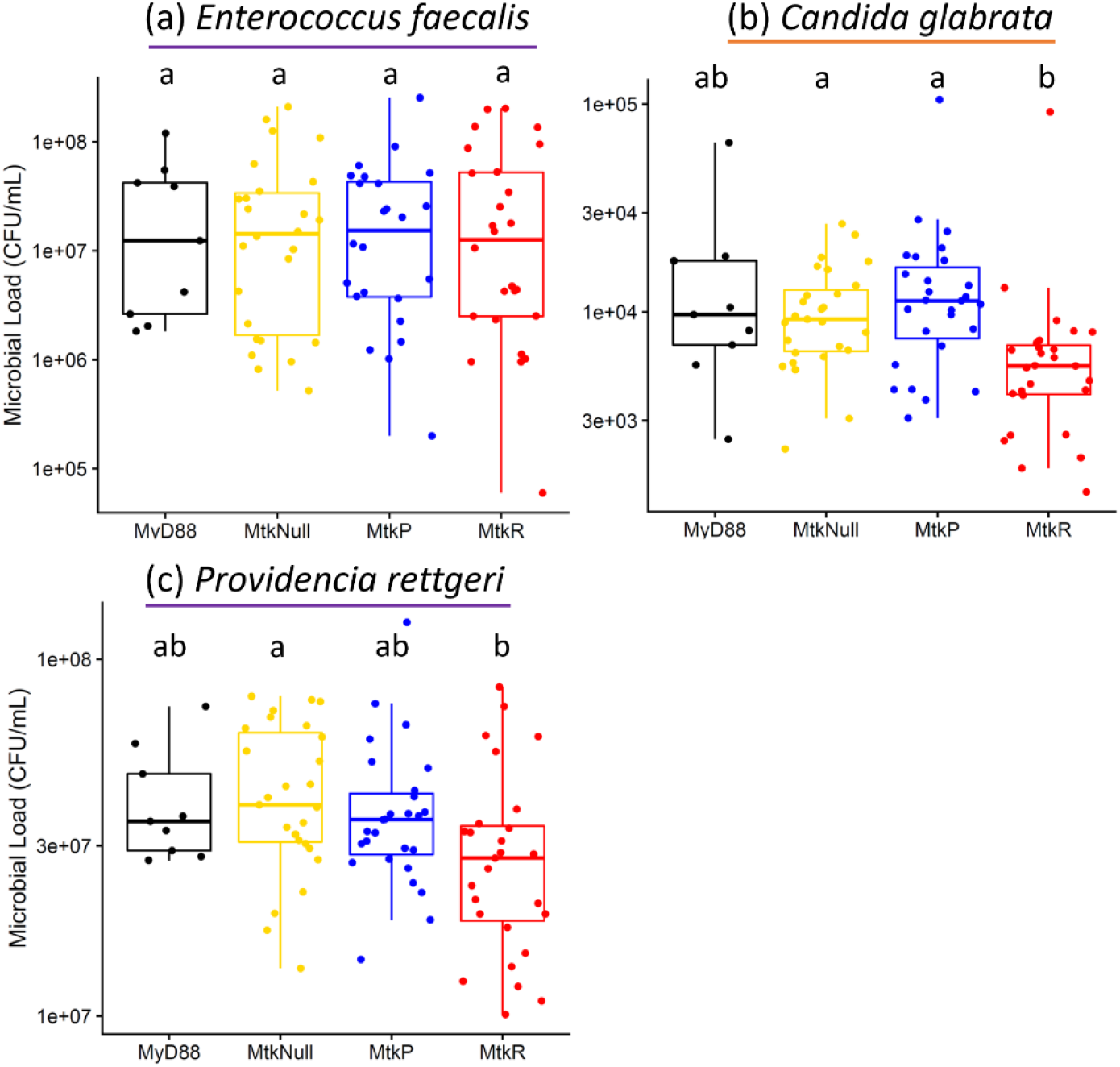
Pathogen load differences suggest that greater pathogen resistance may underlie the greater survival of MtkR males with certain infections. Infections were performed in males with the indicated microbes, using bacteria (purple underline) or yeast (orange underline). Each dot represents the pathogen load 24 hours after infection for 3 pooled flies. Each set of data represents three independent experiments combined. The boxes indicate the interquartile range. Outer edges of the box indicate 25^th^ (lower) and 75^th^ (upper) percentiles and the middle line indicates 50^th^ percentile (median). Whiskers represent maximum and minimum ranges of data within 1.5 times the interquartile range of the box. Letters indicate statistical significance groups, based on a logistic regression and Tukey post hoc test (Table S1).

### *dnaK* is unlikely to play a role in allelic differences in survival

*dnaK* has been previously implicated as a potential microbial target for both Mtk and another proline-rich AMP^57^ and crystal structures of *dnak* and Metchnikowin indicate they bind together (PDB ID 4EZS)^58^. To assess the role of *dnak* in fly infections, we infected our CRISPR-edited *Mtk* lines with a *dnaK* deletion strain of *E. faecalis* to assess if loss of this putative target would result in differential survival among flies of different *Mtk* alleles. However, whether infected with the deletion strain or the parent strain (intact *dnaK*), survival was no different across alleles (Figure S9).

## Discussion

Animals must maintain a delicate balance in their immune systems to maximize survival during infection while reducing any off-target damage to themselves. Prior work in *Drosophila* has indicated that AMPs in particular, compared to all other immune genes, are under balancing selection. Hypotheses suggest that maintenance of allelic variation may relate to this balance between host fitness and antimicrobial activity^31^. Indeed, several specific AMPs have been identified that are likely maintained as multiple alleles in fly populations across the globe despite expectations of rapid evolution^29,30^. However, there have been few investigations either empirically validating allelic functional differences or identifying potential mechanisms underlying this balancing selection^29,31^. The two main hypotheses for balancing selection include the autoimmune hypothesis (one allele is costly without infection, but beneficial with infection) and the specificity hypothesis (each allele is stronger against a different suite of pathogens). Here, with CRISPR/Cas9-edited fly strains, we show strong in vivo evidence of functional differences between two AMPs that differ by only a single amino acid. We find support broadly for the autoimmune hypothesis, in that the MtkR peptide exhibits greater antimicrobial activity than its counterparts in many contexts while simultaneously carrying a host fitness cost. However, when incorporating host sex, we find a more complex picture where sex interacts with host genotype in predicting survival after infection with some pathogens. Broadly, the results support a model where host genotype, sex, pathogen species, and infection status collectively and differentially impact host survival outcomes.

The life history assay revealed several points of interest regarding the relative roles of the *Mtk* alleles in flies (Figure 1). First, egg hatch was lower in flies with the *MtkR* allele. This fitness cost would support the autoimmune hypothesis. Since lower egg hatch occurred in the absence of infection and during the embryonic development stage, this may indicate a role of *Mtk* outside of pathogen response in a process that is important in early development. Indeed, the Toll pathway is known to play an important role in embryonic segmentation^59^. Or, it could indicate leaky *Mtk* expression that has off-target effects in the host. In fact, there is evidence for *Mtk* expression in late-stage embryos^37,60^. Second, there were differences in longevity among alleles and between the sexes. For example, the *MtkR* allele was the costliest in both sexes, but more so in females. Also, the *MtkNull* flies lived the longest with both sexes, but the difference was bigger in males. These results have a few potential implications for *Mtk* function. For one, this indicates that both alleles carry a cost even without infection and regardless of sex. In addition, this cost in the absence of infection coupled with lower egg hatch results further indicates a potential role of *Mtk* in processes outside of pathogen responses that may be important in both embryos and adults. Alternative, leaky expression could be harmful to both developmental stages. Also, the fact that the difference in survival between the two alleles and *MtkNull* flies is larger in males may indicate that the *Mtk* alleles have an important sex-biased function in females or that a putative off-target effect that is particularly harmful to males. Finally, the fact that *MtkR* has the most significantly reduced longevity in the absence of infection along with lower egg hatch rates again supports the autoimmune hypothesis. The reasons for these sex differences are unclear and will need further study. For example, they could potentially be due to differences in food consumption, where females might eat more due to their high reproductive output and incidentally encounter more pathogens and require *Mtk* more often. Or, sex differences in the results could be due to a role of *Mtk* in immune responses to mating or reproduction^47^, or other functions in the host unelated to pathogen responses that may differ between sexes such as microbiome regulation. Specifically, it is possible that *MtkR* may have a more negative interaction with the gut microbiome than *MtkP*, so *MtkR* may lead to dysbiosis in females more so than males. But females, may depend on the Mtk peptides more to offset pathogen encounters with greater feeding. This would be supported by *MtkNull* flies having higher survival in both sexes, but with a greater difference in males. Indeed, females are known to feed more than males^61^ and sex is an important factor that interacts with the fly microbiome to influence factors like host gene transcription and behavior^62,63^.

Systemic infections in males also revealed some interesting patterns. Notably, *MtkR* flies outlive the *MtkP* flies with some infections (Figures 2,3). In other cases, both alleles resulted in similarly higher survival compared to *MtkNull* flies, indicating that any *Mtk* allele is important in these infections. In still other cases, and unexpectedly, both the *MtkR* and *MtkNull* flies had higher survival than *MtkP* flies. In this last case, we imagine that functional Mtk might interact with the pathogen in some complex way to enhance mortality. This may be through interaction with pleiotropic functions of Mtk in the brain^46-48^, or through negative regulatory or direct interactions with secreted fungal molecules, which can occur with Mtk and other AMPs^64,65^. This suggests that host outcomes depend on a complex set of factors including pathogen identity and host genotype and the exact mechanism of functional differences may not simply be one allele being more potent against microbes.

Concerning pathogen species, male infections indicate that a wide variety of fungi and bacteria are inhibited by *Mtk* (Figures 2,3). Expanding on prior results, we show that *Mtk* expression is important in defense against some filamentous fungal, yeast, and bacterial infections. Notably, others have shown that *Beauveria bassiana*^37^, *Micrococcus luteus*^38^, *Neurospora crassa*^37^, and *Fusarium graminaerum*^45^, are inhibited by *Mtk*. Our results support a role of *Mtk* in *Beauveria bassiana* infection and expand on previous findings by demonstrating allelic differences in survival. We find additional evidence that other microbes are inhibited by this AMP: filamentous fungi *Aspergillus fumigatus, Aspergillus flavus*, and *Fusarium oxysporum*, yeasts *Candida glabrata* and *Galactomyces pseudocandidus*, and bacteria *Bacillus thuringiensis* and *Providencia rettgeri*. Thus, there is agreement with prior work on the role of *Mtk* in fighting a broad range of microbial infections, and agreement at the genus level as well. Notably, we did not find a role of *Mtk* in *Candida albicans* infection as previously reported^39^, however, this is likely due to a lower pathogen inoculum here or differences in yeast strains. Importantly, both *Mtk* alleles had higher survival with *Candida albicans* compared to *MtkNull* flies in this study (Figure S6b), but they were not statistically significant with these conditions. We also find no evidence of a role of *Mtk* in infections with *Enterococcus faecalis, Serratia marcescens, Lysinibacillus fusiformis, Staphylococcus succinus, Staphylococcus sciuri, Lactococcus brevis*, and *Lactococcus plantarum*. Thus, we find a few patterns. One is that *Mtk* exhibits activity against most filamentous fungi tested, several yeasts, and only a small subset of mostly Gram-positive bacteria. Another is that while it active against gram-positive bacteria, as previous studies have indicated^38^, there may be some activity against Gram-negative bacteria as well. A final point is that many of the microbes that are not inhibited by *Mtk* include members of the host microbiome (*Lactococcus* species in particular were isolated from flies). There are many possible reasons for this, including *Mtk* being selectively induced against pathogens to avoid disturbing the host microbiome or that these strains are simply not as pathogenic as the others and do not elicit strong enough immune responses to reveal differences in alleles. Indeed, Mtk is expressed in the trachea, surface epithelia, and gut in addition to the fat body^41,42^, so it is possible that normal flora are not impacted by Mtk, or are differentially impacted.

Regarding female infections compared to males, we find many interesting patterns as well (Figures 4,5). In some cases, male and female results are similar such as with *Candida glabrata* (where both alleles help fight infection) and *E. faecalis* (no role of *Mtk*). In other cases, the *Mtk* allele matters in males (*P. rettgeri* and *A. fumigatus*), but there is no role of *Mtk* for females. Finally, we have one case, *B. bassiana*, where *MtkP* is beneficial and *MtkR* is detrimental in females, while the opposite is true of males. It is still unclear why these differences may exist, although it does suggest that *Mtk*, both in general and broken down by allele, can have remarkably different functions in both sexes that significantly affect survival and fitness. Future work will be necessary to determine the mechanism behind sex-based differences. For example, differential regulation of alleles may occur due to reproductive-immune tradeoff, or Mtk may regulate the microbiome in functionally important and sex-dependent ways. Future work will also be required to determine why in some cases, one allele not only performs worse than the other, but also worse even than *MtkNull*. The case of *B. bassiana* is particularly interesting, as the effects in males and females are flipped. Notably, previous work has demonstrated a sexual dimorphism in *D. melanogaster* survival with *B. bassiana*, where females are more susceptible due to differences in immune pathway signaling^66^. Loss-of-function mutations in Toll pathway or *relish* (an Imd gene) removes the dimorphism^66^. Our work expands the finding of *B. bassiana* sexual dimorphism to include a *Mtk* allelic dependency. However, it is still unclear why either allele can be actively detrimental in certain conditions, and future work will be needed to assess this further.

We also took initial steps into assessing the mechanism of the generally more potent *MtkR* activity and found that microbial suppression likely underlies it (Figure 6, Table 1). There is a lower microbial load in *MtkR* flies with *C. glabrata* and *P. rettgeri*. In contrast, *E. faecalis*, which was not impacted by *Mtk* allele, showed no difference in pathogen load. These results mostly parallel the infection survival assays and indicate that *MtkR* is either actively killing or inhibiting growth of the pathogens as a part of its mechanism. In the future, it will be important to understand why. Is this due to expression level differences or differences in activity against the microbial target? In addition, evidence does not support *dnak* being an important microbial target, as with or without the gene, *E. faecalis* infection results in similar survival rates for alleles. Some points remain unclear. For example, we do not know the microbial target of Mtk beyond a doubt. Previous studies to identify a target are based on experiments in a heterologous yeast system with filamentous fungal targets and the results have not been confirmed in a natural system or through other methods^44,45^. Mtk may also have different mechanisms of antimicrobial activity in fungi vs bacteria, for example. This indicates there are many as-yet-undetermined host factors that impact survival regardless of *Mtk* allele.

Here we present evidence of several new aspects of Mtk biology: 1) *Mtk* alleles are functionally distinct. 2) Differential fitness in the context of host allele, host sex, pathogen, and infection status is likely to underlie *Mtk* allele maintenance in wild fly populations. This is because in any particular context, one allele or the other may give the host greater fitness. 2) Differences in host survival can be based on even a single amino acid polymorphism between hosts. 3) New pathogen species in addition to those from previous studies are impacted by *Mtk* expression. 4) Host sex plays a critical role in the outcome of infection with various alleles. 5) We find support through *Mtk* of the immune hypothesis, as broadly speaking, *MtkR* appears to be more potent in many infectious contexts with a fitness cost in the absence of infection. The story is complicated with additional factors like host sex, as either allele may outperform the other depending on sex and pathogen species. This supports the specificity hypothesis to an extent. 6) Finally, while expanding our understanding of *Mtk* function and population-level differences, we also indicate several important avenues of future research. Most notably, further work will elucidate the sex-based differences in *Mtk* allele outcomes along with other aspects of *Mtk* biology, such as its microbial target and putative non-pathogen response host functions.

## Materials and Methods

### Generation and validation of CRISPR fly strains

An inbred *w*^*1118*^ stock was used as the wildtype line for CRISPR/Cas9 genome editing. As such, the strategy was to inject Cas9 protein, guide RNA (gRNA), and single-stranded donor DNA containing the appropriate edits. Lines were created with the intended edit (a single nucleotide mutation resulting in a change from proline to arginine in the 24^th^ residue of the mature Mtk peptide) via homology directed repair, null alleles resulting from nonhomologous end joining, and unedited *Mtk* alleles carried by flies that had been subject to the same injections as the edits and the nulls (Figure S1). Two gRNAs (Mtk_gRNA_target6 and Mtk_gRNA_target14) were designed with PAM sequences just upstream of our desired edit site and synthesized them using the New England Biolabs EnGen sgRNA Synthesis Kit, *S. pyogenes* Protocol (NEB #3322, New England Biolabs, Ipswich, MA, USA). A 120 bp single stranded donor oligo (Mtk_ssDNA1) was used with the desired edit (changing the proline codon – CCA – to an arginine codon – CGA, as well as editing the PAM site to protect against further edits) (Eurofins, Fisher Scientific LLC, Chicago, IL, USA). The gRNAs, single stranded donor oligo, and Cas9 (PNA Biosciences Cas9 Protein with NLS, NC1279639, Fisher Scientific, LLC, Chicago, IL, USA) were injected into 240 embryos (240 separate embryos per gRNA) by GenetiVision Corporation (Houston, TX, USA).

Several of the injected embryos did not survive to adulthood, but those that did were mated to the parent *w*^*1118*^ line and their offspring were Sanger sequenced using primers Mtk_F1 and Mtk_R1 (Table S2) as a brute force approach to finding edits. Any F1 individuals with evidence of edits (double peaks that begin around the edit site) were inbred to isolate and create homozygous mutants. Siblings of our putative edited individuals were kept as unedited controls. Through several crosses and Sanger sequencing, homozygous lines were created with our edit (3 lines that are likely 3 different edits since they were derived from different G0 embryos) or deletions (1bp, 6bp or 17bp), as well as 3 unedited control lines (Figure S1).

To ensure that these lines did not significantly differ in sequence other than the focal edits in *Mtk*, light whole genome sequencing was performed on each of the nine lines derived from *w*^*1118*^. Briefly, DNA was extracted from each line using the Qiagen Gentra Puregene Tissue Kit (158066, Qiagen, Germantown, MD, USA), then prepared library using the Illumina Nextera DNA Library Preparation Kit (Illumina, Inc., San Diego, CA, USA). Samples were sequenced on a fraction of a HiSeq2000 lane to an average depth ranging from 4 to 12. As a control, three DGRP (391, 405 and 280) lines were also used. As expected, the number of pairwise differences between any of our CRISPR/Cas9 genome edited lines was low (less than 200 putative differences per chromosome). In comparison the number of pairwise differences per chromosome between any of our lines and the DGRP lines ranged from 40,000 to 80,000. So, while the possibility that the editing approach caused off target effects in other essential genes cannot be ruled out, there is no evidence that this is the case, and our expectation is that most of the SNP differences between CRISPR/Cas9 edited lines are either mapping artifacts or the product of standard mutational processes.

### Fly strains and husbandry

Additional fly strains include *spz* (*spz/rm7*, loss of function point mutation^67^, gift from B. Lazzaro), and *MyD88* (BDSC 14091, transposable element insertion). Flies were reared on CMY media: 64.3 g/L cornmeal (Flystuff Genesee Scientific, San Diego CA), 79.7 mL/L molasses (Flystuff Genesee Scientific, San Diego CA), 35.9 g/L yeast (Genesee Scientific, San Diego CA, inactive dry yeast nutritional flakes), 8 g/L agar (Flystuff Genesee Scientific, San Diego CA, *Drosophila* type II agar), 15.4 mL of antimicrobial mixture [50 mL phosphoric acid (Thermo Fisher, Waltham MA), 418 mL propionic acid (Thermo Fisher, Waltham MA), and 532 mL deionized water], and 1g/L tegosept (Genesee Scientific, San Diego CA). Flies were reared and collected at room temperature (25°C) unless otherwise stated and were kept on a 16h light/8 h dark light cycle.

### Microbial strains and growth conditions for fly infections

The following microorganisms were used in this study: *Providencia rettgeri* Dmel (gram negative isolate from wild *D. melanogaster* by B. Lazzaro in State College, PA, USA), *Bacillus thuringiensis* Berliner (gram positive isolate strain ATCC 10792 isolated from *Ephestia kuehniella*), *Enterococcus faecalis* Dmel (gram positive isolate from wild *D. melanogaster* by B. Lazzaro in State College, PA, USA), *Serratia marcescens* (gram negative isolate by R. Unckless from *D. melanogaster* hemolymph), *Lysinibacillus fusiformis* (gram positive strain Juneji, isolate from *D. melanogaster*, gift from B. Lazzaro), *Staphylococcus succinus* (gram positive, isolate from *Drosophila* by R. Unckless), *Staphylococcus sciuri* (gram positive, isolate from *Drosophila* by R. Unckless), *Lactococcus brevis* (gram positive), *Lactococcus plantarum* (gram positive), *Enterococcus faecalis* (gram positive strain K-12 CGSC 7636, isolated from feces), *Enterococcus faecalis* (strain K-12 ΔdnaK CGSC 8342, derivative of CGSC 7636), *Candida glabrata* strain CBS 138 (yeast strain ATCC2001, isolated from feces), *Galactomyces pseudocandidus* (yeast, isolated from contaminated *Drosophila* stock by I. Nevarez-Saenz), *Candida auris* (yeast strain CDC B11903, clinical isolate), *Candida albicans* (yeast strain SC5314, clinical isolate), *Fusarium oxysporum* f. sp. Lycopersici (Fungal Genetics Stock Center strain 9935, isolated from tomatoes), *Beauveria bassiana* strain GHA (gift from P. Shahrestani, isolated from *Locusta migratoria*), *Aspergillus fumigatus* (Fungal Genetics Stock Center strain 1100, clinical isolate), and *Aspergillus flavus* strain NRRL 3357 (Fungal Genetics Stock Center strain A1446, isolated from peanuts).

To grow cultures for fly infections, bacteria and yeast isolates were grown overnight for 16 h from a single colony in 2 mL media with shaking at 225 rpm. Most bacteria were grown in Luria broth (LB) (Teknova, Hollister CA) from colonies on Luria agar (LA) (Teknova, Hollister CA, molecular biology grade agar) at 37°C and yeast were grown in potato dextrose broth (PDB, BD, Sparks MA) from colonies on potato dextrose agar (PDA) at 30°C. *Lactococcus* species were grown in MRS broth (BD, Sparks MA) from colonies on MRS agar plates at 30°C. Isolates were then prepared as described below. Fungal spores were prepared by purifying spores grown on PDA at 30°C for 1-2 weeks. Autoclaved DI water was poured over each plate and the spores were suspended in the liquid. This was then poured over a filter (Millipore Sigma, Burlington MA, Miracloth 22-25 µm pore size) and the filtrate was placed into a 50 mL falcon tube. This was then centrifuged at 1000 rpm for 5 min and the supernatant was discarded. The spores were then resuspended in sterile 20% glycerol and were counted using a hemocytometer and stored at 4°C until use.

### Fly infections

Bacteria and yeast were grown overnight in the conditions described above. Yeasts *C. glabrata, C. auris, C. albicans*, and *G. pseudocandidus* were diluted in PDB to an optical density (OD) value of A_600_=35 +/-1 for *Candida glabrata* and *Candida auris*, an OD value of A_600_=250 +/-5 for *Candida albicans*, and an OD value of A_600_= 120+/-1 for *Galactomyces*. Bacteria were diluted in LB to and an OD value of A_600_= 4 +/-0.02 for *B. thuringiensis, E. faecalis, L. brevis, L. plantarum*, and *S. marcescens*, an OD value of A_600_= 1.5 +/-0.02 for *E. faecalis* K-12 and Δ*dnaK*, and an OD value of A_600_= 1 +/-0.02 for *P. rettgeri*. Sporulating fungi were prepared as described above. Males or females 4-6 days old of a given genotype were pierced in the thorax just beneath the wing using a 0.15 mm dissecting pin (Entosphinx, Czech Republic, No. 15 Minuten pins 12 mm long 0.15 mm diameter) dipped into the diluted culture or control. Controls were the growth broth for yeasts and bacteria (PDB or LB, see above) or sterile 20% glycerol for the fungal spores. Flies were then placed in groups of 10 per food vial. 20 individuals of each treatment x sex x isoline group were infected in each block (3 isolines per genotype), and at least two blocks of infections were performed on separate days for every experiment. Flies infected with fungi were counted daily for survival for 21 days and those infected with bacteria were counted for 7 days, as differences in survival across treatment groups were observed within these time periods.

### Life history assay

Virgin females of each isoline of the *Mtk* CRISPR flies were collected and aged 2-4 days. They were then crossed to males of the same isoline, also 2-4 days old. This was done by placing single male-female pairs of each isoline into a 6 oz. square bottom *Drosophila* bottle (Fisher Scientific, Hampton NH) covered with a grape juice agar plate [100% concord grape juice (Welch’s, MA), tegosept (Genesee Scientific, San Diego CA), 200-proof ethanol (Decon Laboratories Inc, PA), agar (Teknova, Hollister CA), DI water] with yeast paste (Fleischmann’s Active Dry Yeast, Heilsbronn Germany). There were 32 crosses per isoline for the first block and 16 crosses per isoline for the second. These bottles were placed in a 25°C incubator overnight (12-hour dark/light cycle). Grape plates were swapped the next morning (16 hr later) with fresh plates and yeast. The bottles were placed back in the incubator and flies were allowed to lay eggs for 24 h. Then, plates were removed and adults discarded. The number of eggs was immediately counted and the plates were placed back into the incubator. The number of hatched embryos was counted the next day and larvae were moved into vials with CMY *Drosophila* media and the vials were kept in the incubator. Both pupae and emerged adult offspring were counted. The adult offspring were separated by sex into groups of 20 and placed in fresh CMY food vials and kept in the incubator. Flies were given fresh food vials every 4-7 days. Surviving fly numbers were recorded regularly until fly death.

### Microbial load assay

Microbes were grown overnight in 2 mL culture vials at 225 rpm. Bacteria *E. faecalis* and *P. rettgeri* were grown in LB at 37°C. Yeast *C. glabrata* was grown in PDB at 30°C. After 16 h growth, the cultures were diluted to the desired OD values: A_600_= 4 +/-0.02 for *E. faecalis*, A_600_= 1 +/-0.02 for *P. rettgeri*, and A_600_=35 +/-1 for *C. glabrata*. Infections were performed as described above, by piercing flies in the thorax with a needle dipped in culture or control. For each isoline x treatment combination, 9 3-4 day old males were infected. The flies were kept at room temperature (25°C) for 24 hr. The flies were allocated into 1.7 mL Eppendorf tubes with 3 groups of 3 flies for each isoline x treatment combination. Then, 150 µL sterile PBS (Corning, Corning NY) was added to each along with a single sterile 2 mm glass bead (Merck, Germany) to each tube. The flies were then homogenized in a bead beater (VWR, Radnor PA, Hard Tissue Homogenizer) for 3 minutes at 1600 rpm. The samples were then placed on ice and plated on either LBA (bacteria or LB controls) or PDA (yeast or PDB controls). They were plated using a Whitley WASP Touch Automated Spiral Plater (Don Whitley Scientific, West Yorkshire, UK) using 50 µL of the homogenate per plate. The plates were then placed in incubators overnight (37°C for LBA plates, 30°C for PDA plates). Plates were counted using a colony counter (Neutec Group Inc., Farmingdale NY, IUL Flash & Go Automated Colony Counter).

### Data and statistical analyses

Data analysis and figure generation were performed in R^68^, using several packages: ggplot2^69^ (version 3.4.0), cowplot^70^ (version 1.1.1), car^71^ (version 3.1.1), SurvMiner^72^ (version 0.4.9), MultComp^73^ (version 1.4.20), ggExtra^74^ (version 0.10.0), and SurvMisc^75^ (version 0.5.6). Dot plots were analyzed with a logistic regression and Tukey post-hoc test. Longevity plots without infection were analyzed using ANOVA and Tukey post-hoc test. Longevity plots with infection were analyzed using a Cox proportional hazard model with MtkP as the reference.

## Data Availability

All data are available upon request.

## Acknowledgments

We would like to thank B. Lazzaro and P. Shahrestani for providing certain fly and microbial strains. We would also like to thank the Bloomington Drosophila Stock Center (BDSC) for providing some fly lines. This work was supported by two National Institutes of Health (NIH) K-INBRE P20 GM103418 postdoctoral awards (one each to JIP and JRC), National Science Foundation (NSF) Postdoctoral Fellowship in Biology (PRFB) DBI 2109772 to JIP, and NIH grants R01 AI139154 and P30-GM110761 to RLU.

## Contributions

JIP, JRC, and RLU conceived and designed experiments. JIP and RLU analyzed the data and wrote the manuscript. JIP and JRC performed fly experiments. JRC and RLU generated and validated the CRISPR lines. MCW and INS contributed to mechanistic experiments. All authors approved of the final version of the manuscript.

## Supporting Information

See Table S1 for statistical outputs for each analysis.

---

**Figure S1.**
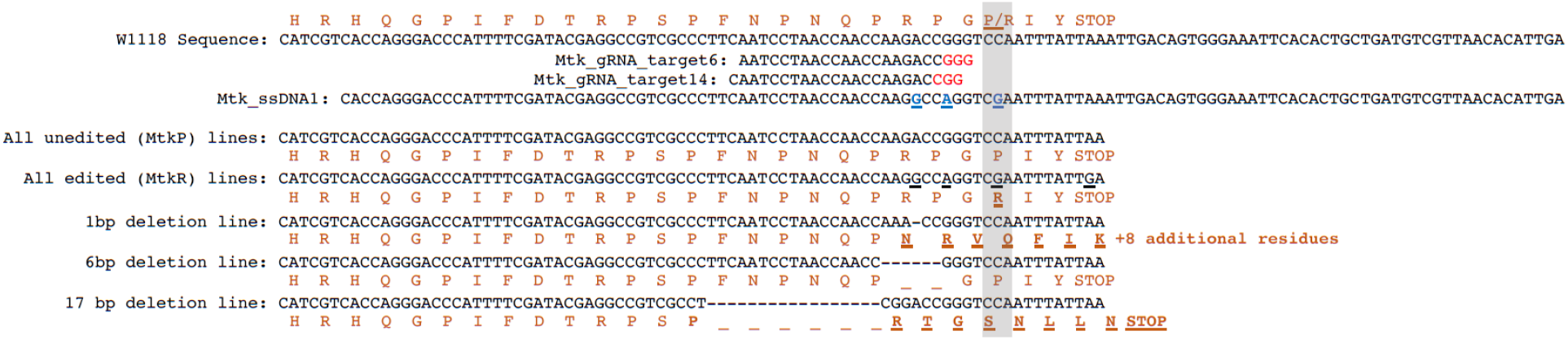
Schematic of CRISPR-generated *Mtk* fly lines. The *w*^*1118*^ sequence represents the starting sequence of the background line (P allele). Two guide RNAs (gRNA) were used to generate flies with the R allele (ssDNA1). The bottom 5 lines depict the nucleotide and amino acid sequences for the lines used in this paper.

**Figure S2.**
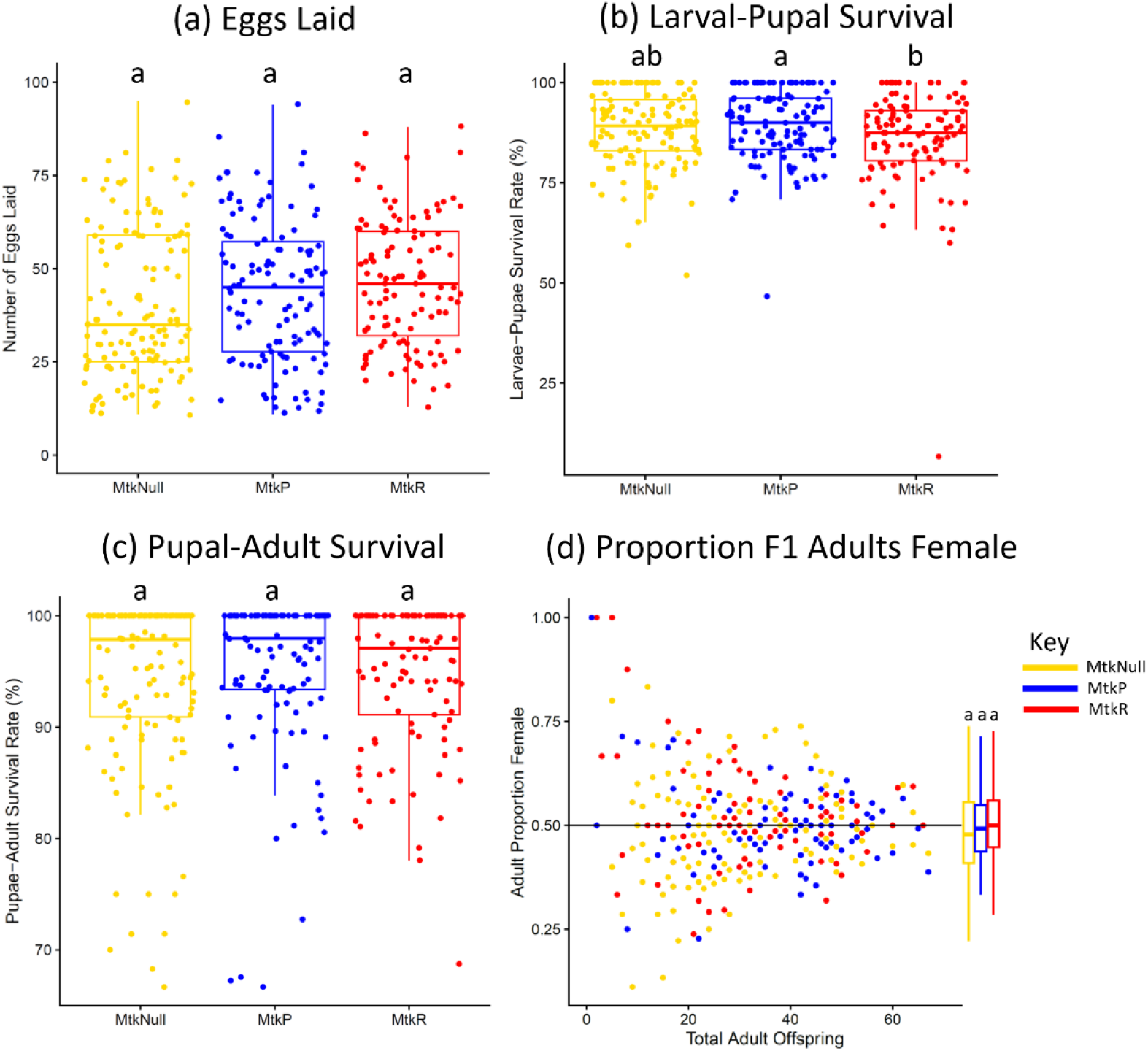
Offspring numbers, survival from larvae to adults, and the proportion of F1 females exhibit small to no differences among *Mtk* alleles. (a) The number of offspring was counted for each family. Each dot represents all eggs from one female (overall mean of 41 eggs). (b) The larvae-to-pupae survival rate was counted for each family. (c) The pupae-to-adult survival rate was counted for each family. (d) The proportion of adult offspring that were female compared to total adult offspring. Outliers are largely due to families with few offspring. Box plots for each genotype are plotted to the right of the scatter plot for comparison among genotypes (no significant differences). For all graphs, each dot represents the offspring of a single male and female of the indicated genotype. The boxes indicate the interquartile range. Outer edges of the box indicate 25^th^ (lower) and 75^th^ (upper) percentiles and the middle line indicates 50^th^ percentile (median). Whiskers represent maximum and minimum ranges of data within 1.5 times the interquartile range of the box. Letters indicate statistical significance groups, based on a logistic regression and Tukey post hoc test (Table S1). Families that laid no eggs were not included (usually due to death of a fly during the experiment; similar number of families excluded across alleles). The entire experiment was performed twice, and graphs represent a combination of data from both experiments.

**Figure S3.**
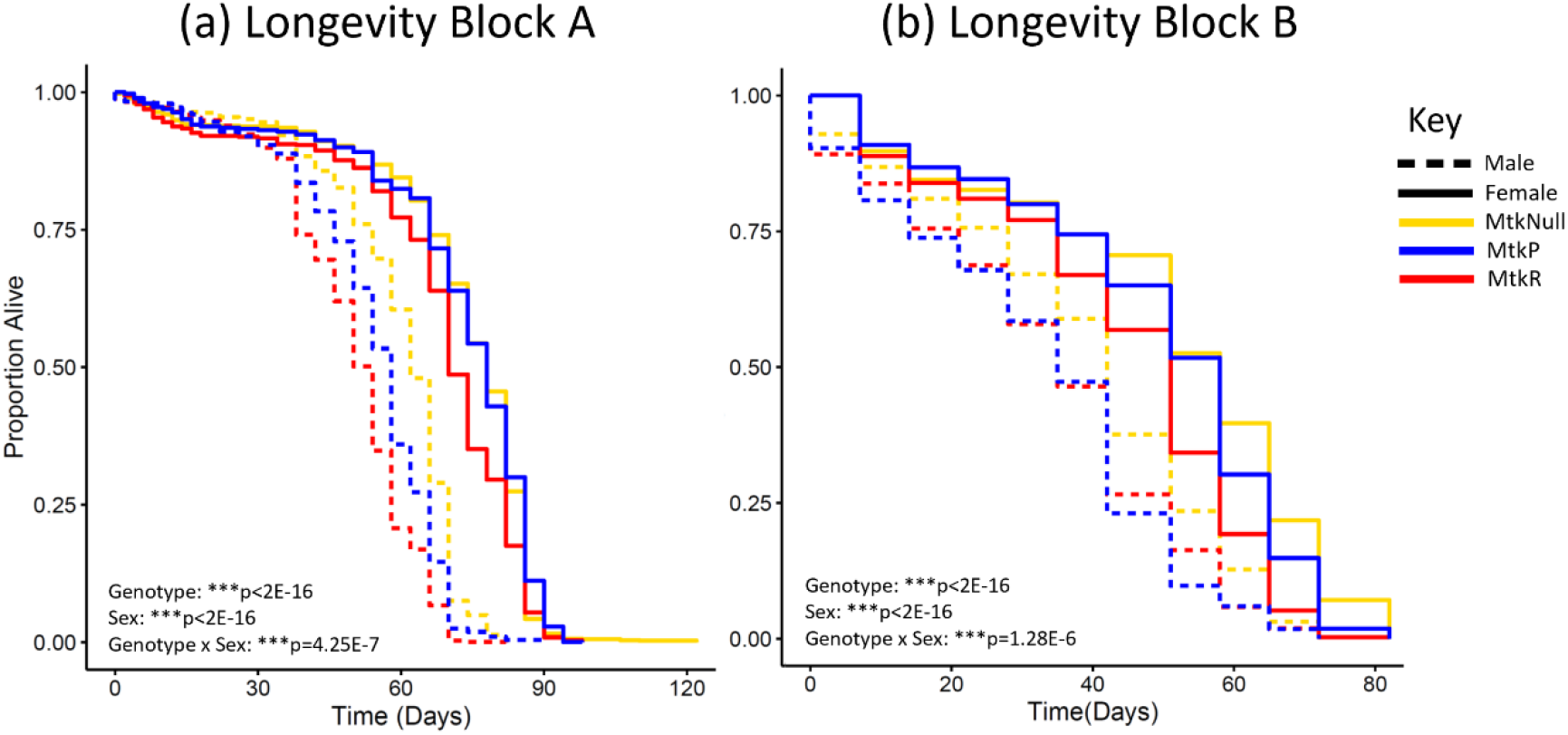
The longevity of adult male *MtkR* varies, but *MtkR* females consistently die earlier. (a) The longevity of adult offspring in block A, with an average of 2182 flies per genotype, sexes combined. (b) The longevity of adult offspring in block B, with an average of 1697 flies per genotype, sexes combined. Statistics based on an ANOVA with Tukey post-hoc test (Table S1). The experiment was performed twice.

**Figure S4.**
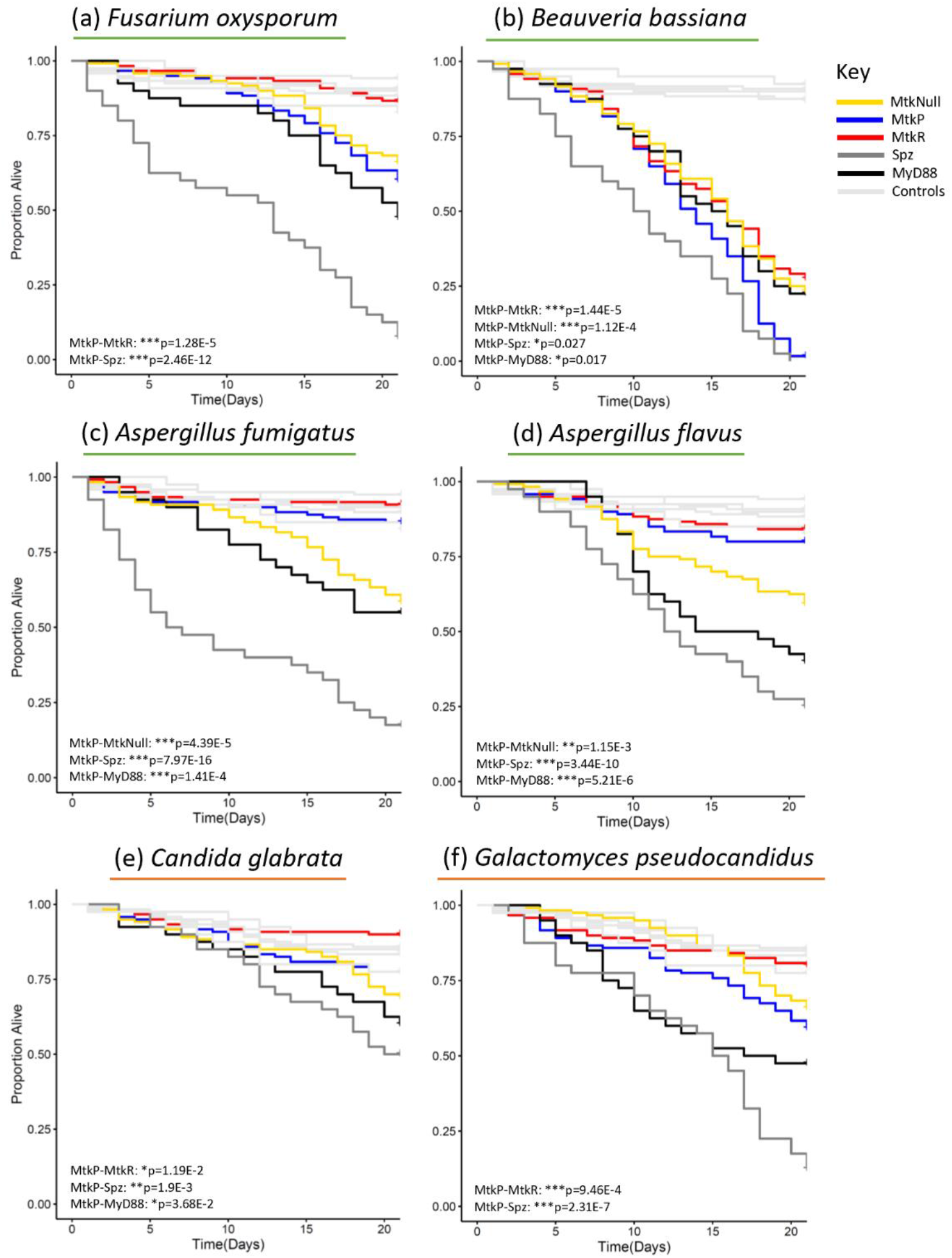
Survival curves for males infected with fungi, corresponding to Figure 2. Infections were performed with the indicated microbes, using either spores (green underline) or yeast cultures (orange underline). Each line represents the survival of 120 flies (*Mtk* alleles and controls) or 40 flies (*Spz* and *MyD88*) over a 21-day period for the same data graphed in Figure 2. Statistics based on cox proportional hazard model (Table S1). The experiment was performed twice, with combined results represented here.

**Figure S5.**
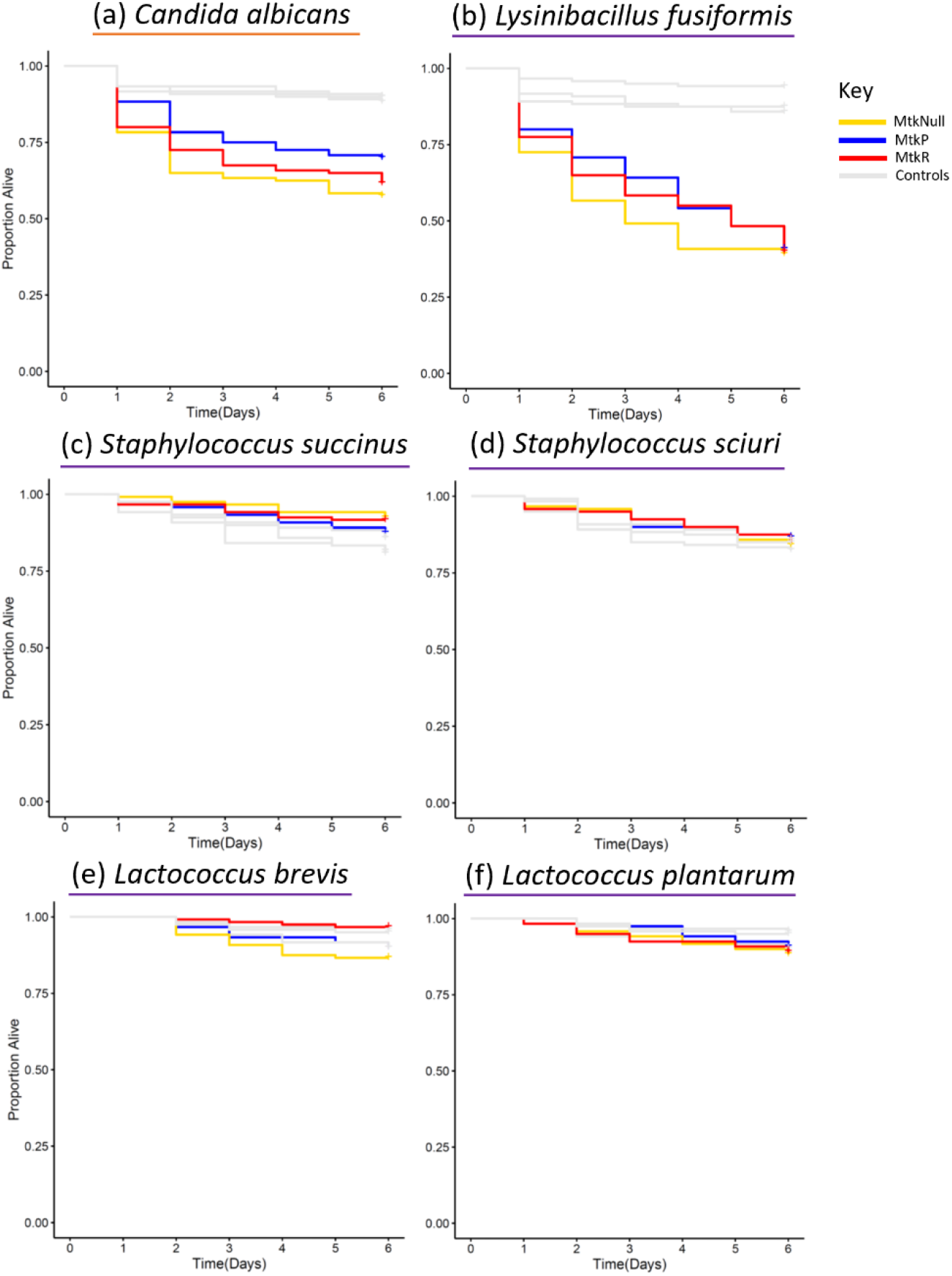
Survival curves for males infected with bacteria or yeast. Infections were performed with the indicated microbes, using bacteria (purple underline) or yeast (orange underline). Each line represents the survival of 120 flies (*Mtk* alleles and controls) or 40 flies (*Spz* and *MyD88*) over a 7-day period. Statistics based on cox proportional hazard model (Table S1). The experiment was performed twice, with combined results represented here.

**Figure S6.**
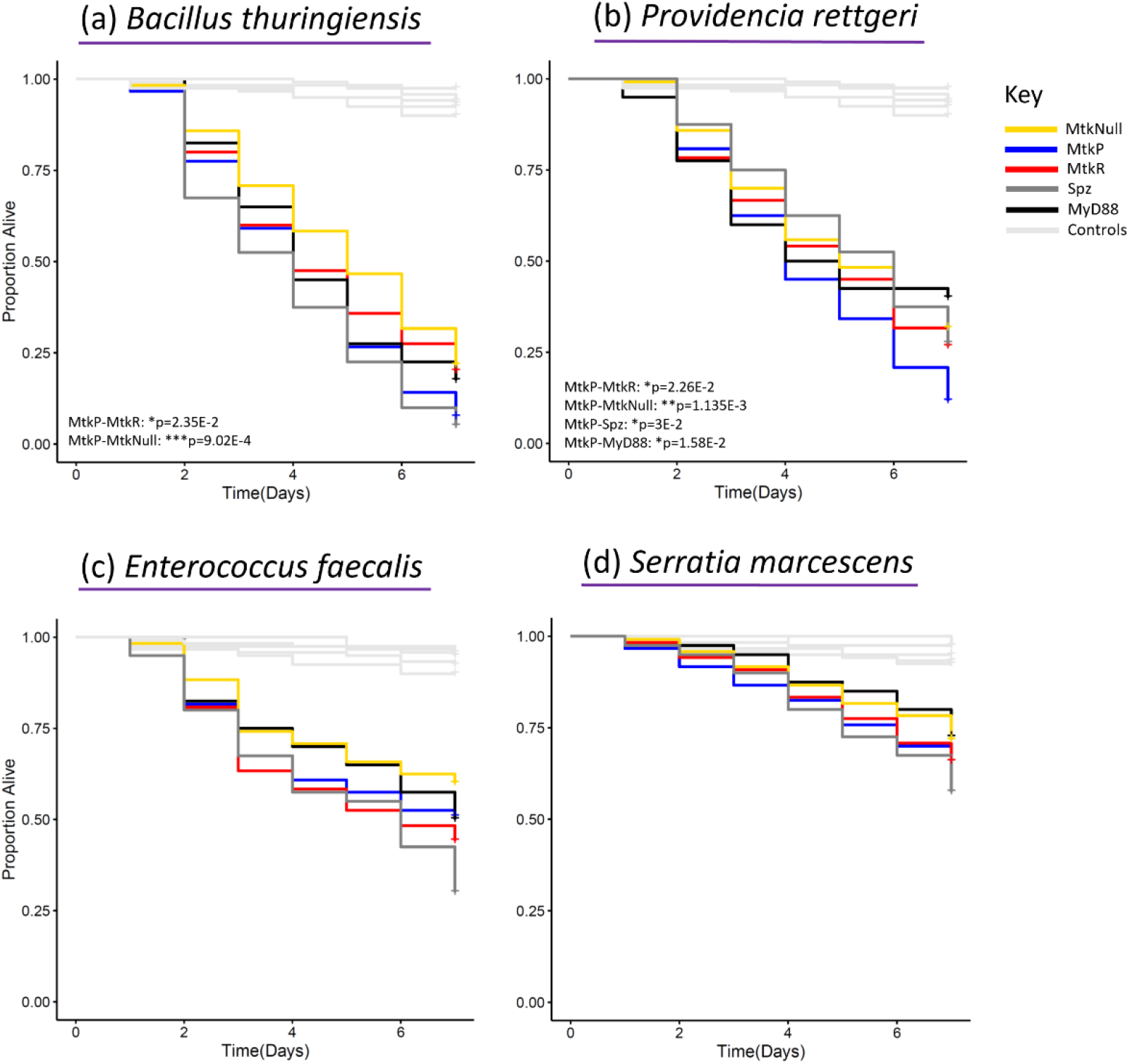
Survival curves for males infected with bacteria, corresponding to Figure 3. Infections were performed with the indicated microbes, using bacteria (purple underline). Each line represents the survival of 120 flies (*Mtk* alleles and controls) or 40 flies (*Spz* and *MyD88*) over a 7-day period for the same data graphed in Figure 3. Statistics based on cox proportional hazard model (Table S1). The experiment was performed twice, with combined results represented here.

**Figure S7.**
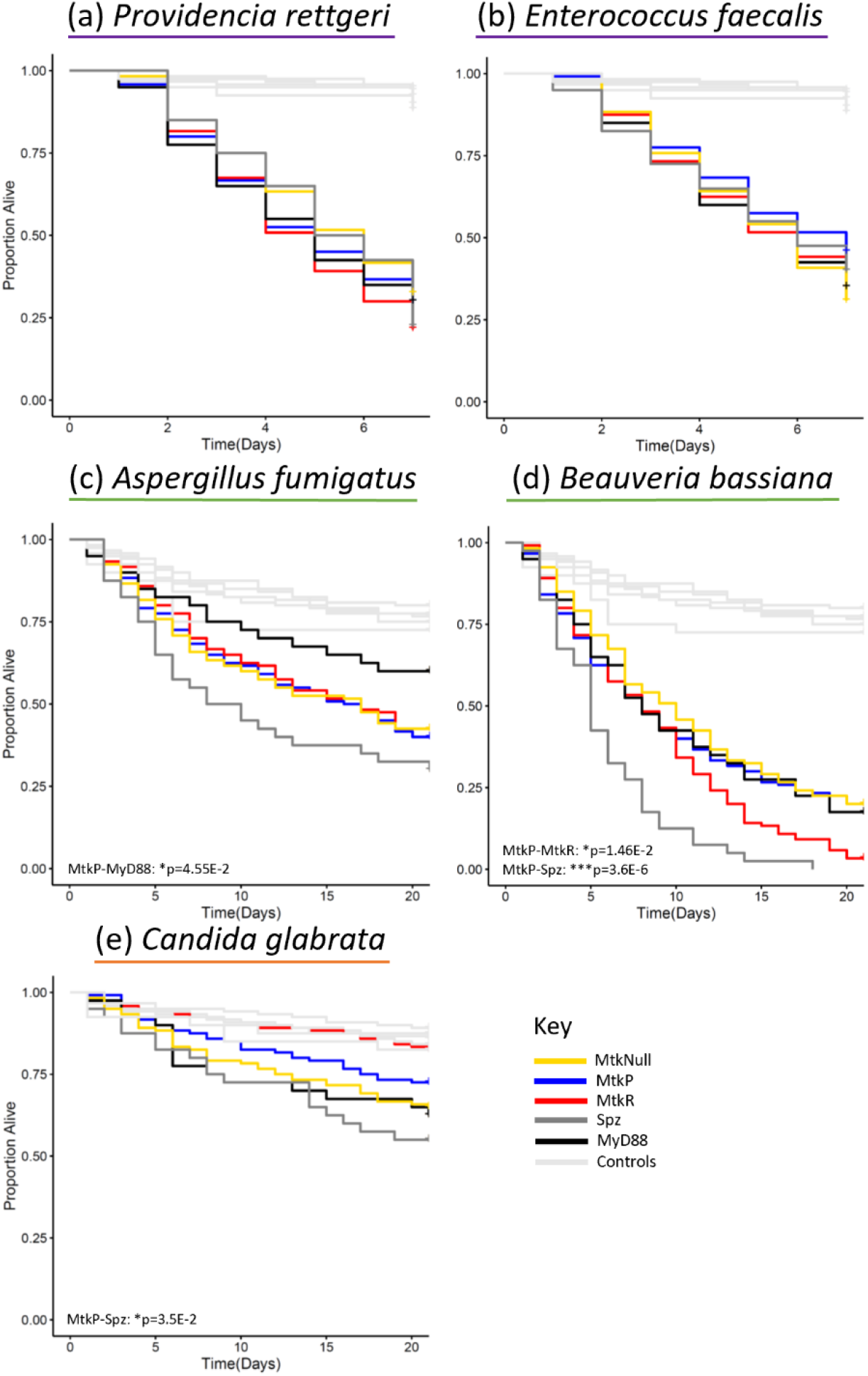
Survival curves for females infected with bacteria, fungal spores, or yeast corresponding to Figure 4. Infections were performed with the indicated microbes, using bacteria (purple underline), spores (green underline), or yeast (orange underline). Each line represents the survival of 120 flies (*Mtk* alleles and controls) or 40 flies (*Spz* and *MyD88*) over a 21-day period. Statistics based on cox proportional hazard model (Table S1). The experiment was performed twice, with combined results represented here.

**Figure S8.**
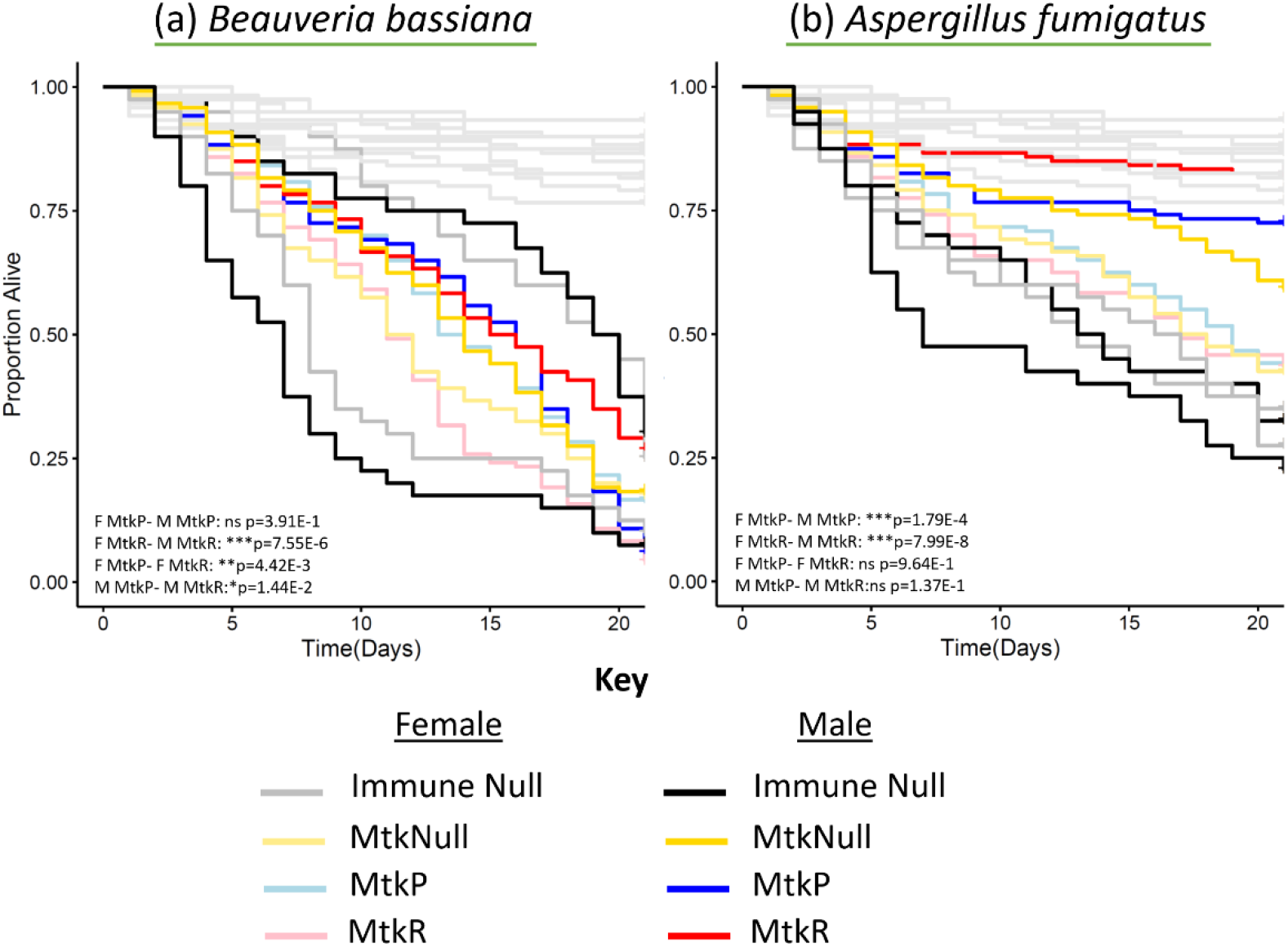
Survival curves for side-by-side male and female fungal spore infections corresponding to Figure 5. Infections were performed with the indicated microbes, using spores (green underline). Each line represents the survival of 120 flies (*Mtk* alleles and controls) or 40 flies (*Spz* and *MyD88*) over a 21-day period. Statistics based on cox proportional hazard model, with FDR for a subset of contrasts (Table S1). The experiment was performed twice, with combined results represented here.

**Figure S9.**
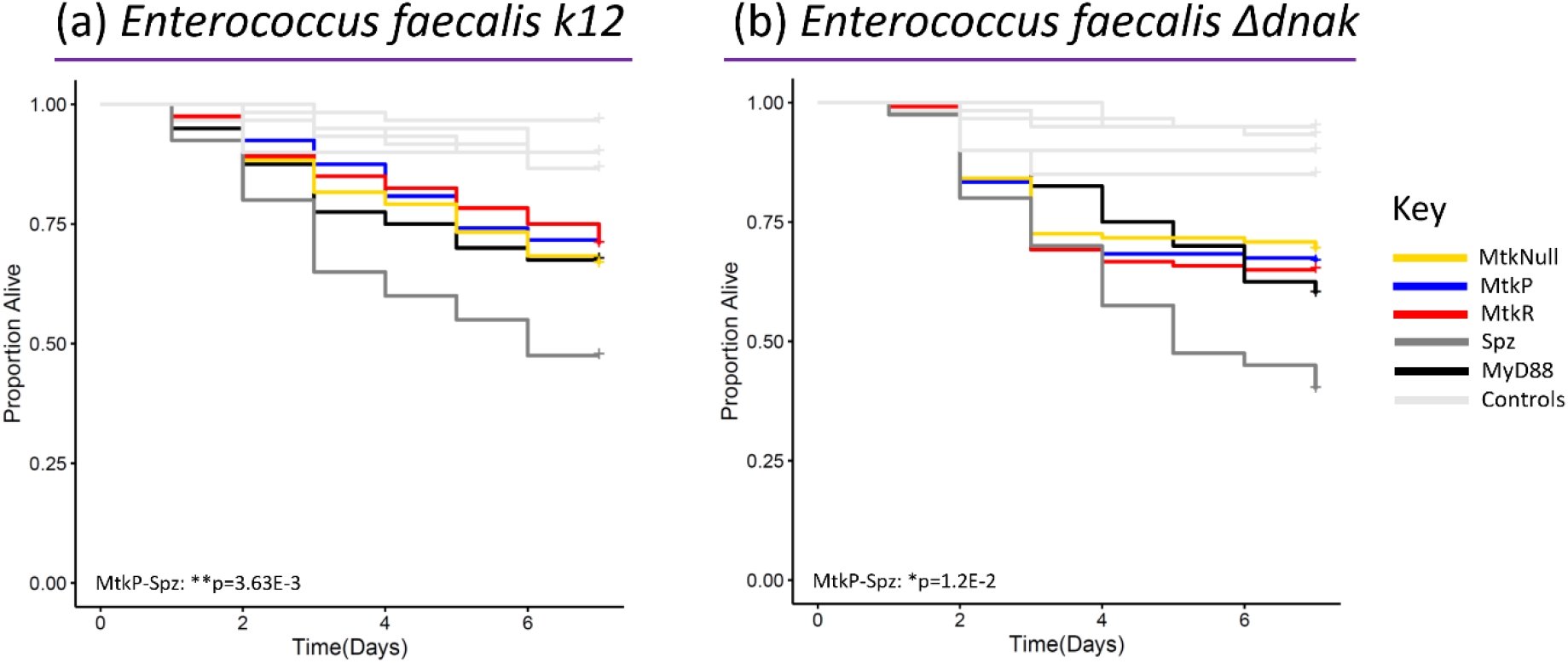
Males infected with *dnaK*-deficient *E. faecalis* (Δ*dnak*) survive at similar rates across alleles. Infections were performed with *E. faecalis* (purple underline). Each line represents the survival of 120 flies (*Mtk* alleles and controls) or 40 flies (*Spz* and *MyD88*) over a 21-day period. Statistics based on cox proportional hazard model (Table S1). The experiment was performed twice, with combined results represented here.

**Table S2.**
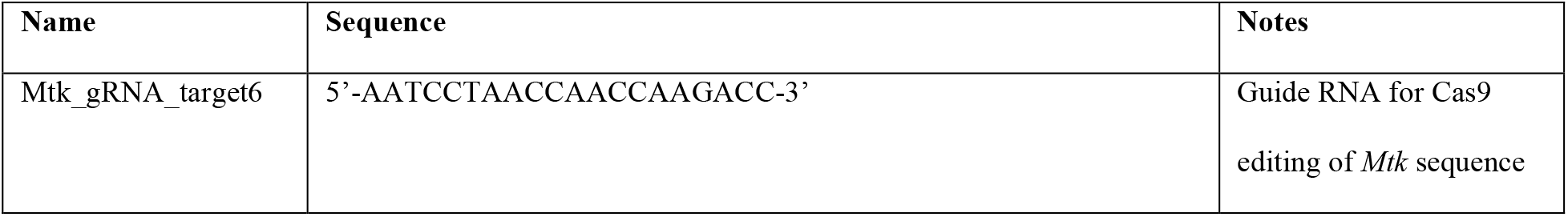
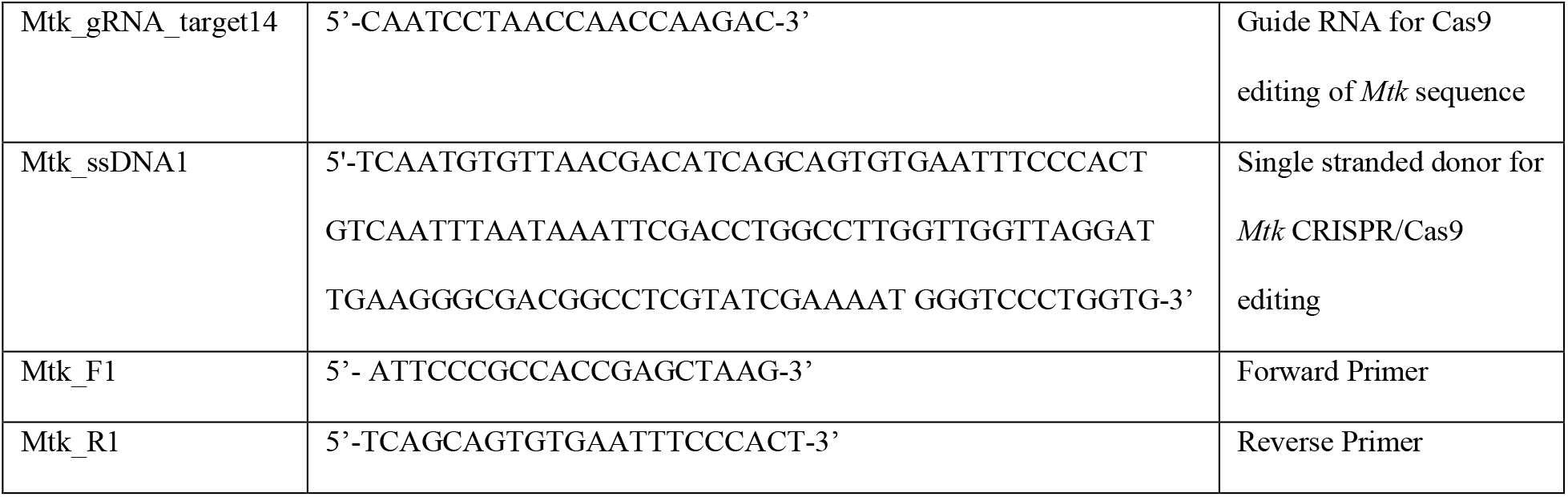
Oligos used in CRISPR/Cas9 editing of *Drosophila Mtk* alleles.

**Table S3.**
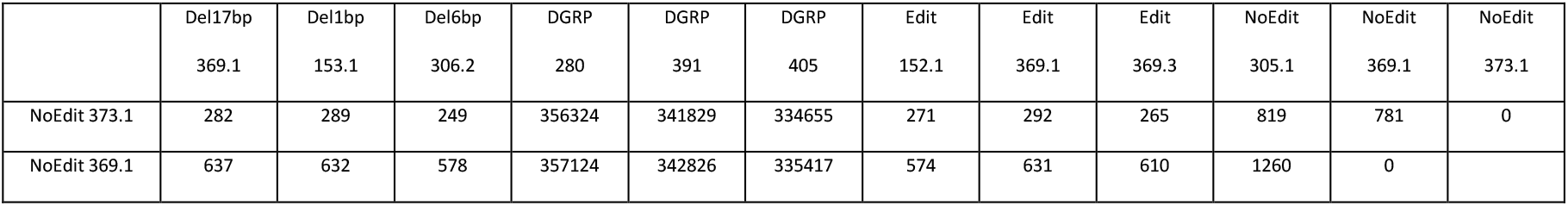
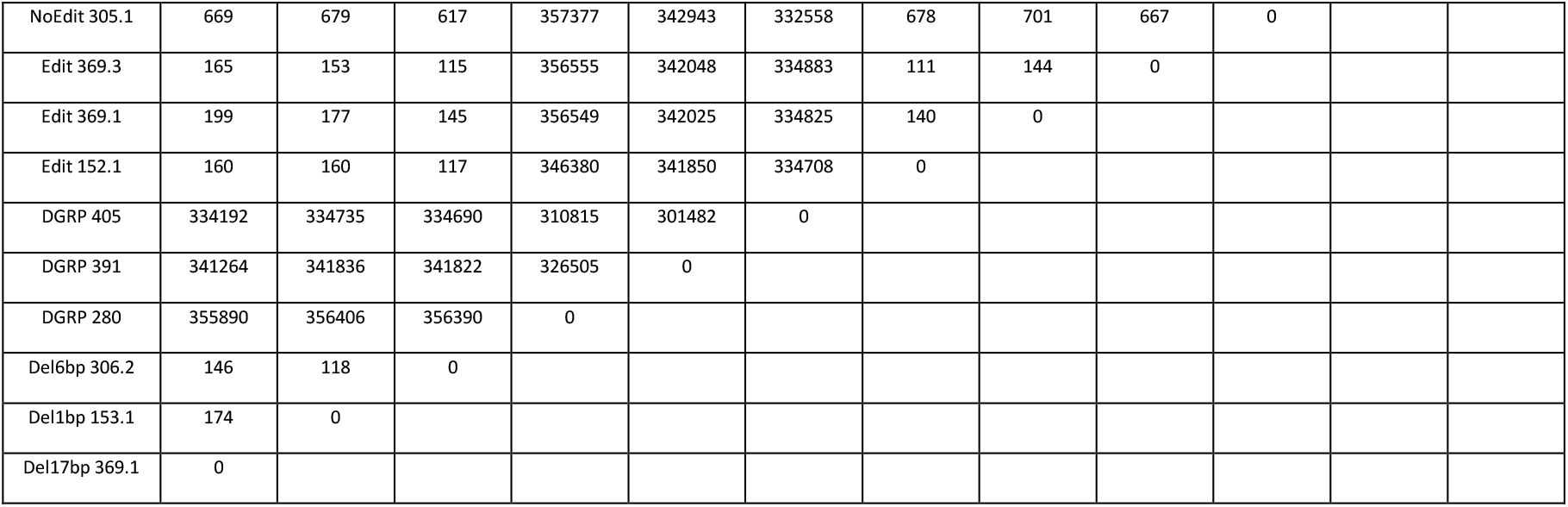
Pairwise SNP differences among whole genomes (2L, 2R, 3L, 3R and X) of CRISPR/Cas9 genome edited lines and three random DGRP lines.

